# Effect of heterozygous deletions on phenotypic changes and dosage compensation in *Arabidopsis thaliana*

**DOI:** 10.1101/2024.09.20.614168

**Authors:** Takuya Ikoma, Ryo Nishijima, Miho Ikeda, Kotaro Ishii, Asanga Deshappriya Nagalla, Tomoko Abe, Yusuke Kazama

## Abstract

Heterozygous deletions, which include a large number of genes, are often caused by the induction of mutations. The induction of gene dosage compensation should be considered when assessing the effects of heterozygous deletions on phenotypic changes. This mechanism is known to balance the expression levels of genes with different copy numbers in sex chromosomes, but it is also known to operate in autosomes. In the present study, 12 *Arabidopsis thaliana* BC_1_ mutants with heterozygous deletions were produced by crossing wild-type Col-0 plants with mutants induced by heavy ion beams. The sizes of the deletions ranged from 50.9 kb to 2.03 Mb, and the number of deleted genes ranged from 8 to 92. Nine of the 12 BC_1_ mutants showed phenotypic changes in fresh weight 14 days after cultivation or during the flowering period. RNA-sequencing (RNA-seq) analyses of 14-day-old leaves, 40-day-old leaves, and flower buds showed that dosage compensation did not occur in any stage or tissue tested. These results indicate that heterozygous deletions cause phenotypic changes owing to the absence of dosage compensation.

**Article Summary:** In this study, we examined the impact of heterozygous deletions in *Arabidopsis thaliana*. By crossing wild-type Col-0 plants with mutants induced by heavy ion beams, we created 12 BC1 mutants, each having heterozygous deletions. The deletions ranged from 50.9 kb to 2.03 Mb, affecting 8 to 92 genes. Nine mutants showed changes in fresh weight or flowering time. RNA-seq analyses of leaves and flower buds revealed no gene dosage compensation. Our findings indicate that phenotypic changes result from the lack of dosage compensation in heterozygous deletions.

## Introduction

Recent advancements in mutation induction, including genome editing (Rönspies et al., 2022), heavy-ion beam irradiation (Ishii et al., 2024), and the Ex-TAQing system (Tanaka et al., 2020), have enabled the induction of largescale mutations, such as translocations and extensive deletions. These techniques often facilitate the generation of heterozygous deletions, primarily to avoid the lethality associated with the homozygous disruption of essential genes (Naito et al., 2005; Ishii et al., 2024). Heterozygous deletions result in an imbalance in gene copy number, which is a critical factor in breeding programmes involving such rearrangements. Therefore, the evaluation of the effects of gene copy number imbalances is essential for successful breeding using these genetic modifications.

An imbalance in gene copy number is sometimes harmful to organisms. For example, aneuploidy is known to be more harmful than polyploidy and results in dynamic phenotypic changes (Blakeslee et al., 1920; Blakeslee, 1921; Blakeslee, 1934; Bridges, 1925; Sinnott and Blakeslee, 1922). Several cancers are associated with trisomy of various chromosomes (Rajagopalan and Lengauer, 2004). This is thought to occur because the balance in the expression of transcription factor genes, high-molecular-weight molecule complexes, or molecules involved in signal transduction is disrupted (Birchler et al., 2005; Birchler and Veitia, 2007; Birchler and Veitia, 2010; Birchler and Veitia, 2012).

Another example of gene imbalance involves sex chromosomes, where the copy numbers of genes differ between males and females or between sex chromosomes and autosomes. To counteract this imbalance, many organisms have evolved a mechanism known as gene dosage compensation to equalise gene expression levels. For instance, in *Drosophila*, the balance of gene expression is maintained by upregulating the expression of X-linked genes in males (Baker et al., 1994; Nozawa et al., 2014; Samata and Akhtar, 2018). In humans, gene expression levels linked to one of the X chromosomes in women are downregulated to balance the gene expression levels (Heard et al., 1997; Disteche, 2012; Graves, 2016). Disruption of gene dosage compensation machinery can lead to developmental defects. The absence of gene dosage compensation in *D. melanogaster* leads to male-specific lethality in third instar larvae (Belote and Lucchesi, 1980).

Sex chromosomes evolve from a pair of autosomes (Furman et al., 2020). Human sex chromosomes diverged from an autosomal pair 180 million years ago (Ohno, 1967; Graves, 1995; Lahn and Page, 1999; Potrzebowski et al., 2008; Veyrunes et al., 2008). Initially, the divergence between the X and Y chromosomes was less pronounced than that in contemporary sex chromosomes. Over time, the Y chromosome degenerated, and the number of X-specific genes increased, resulting in heteromorphic sex chromosomes (Lyon, 1961). Ohno proposed that the upregulation of X-linked genes in early sex chromosomes in males was followed by X chromosome inactivation in females, leading to the current state (Ohno, 1967). In other words, mammalian sex chromosomes also exhibit upregulation of X-linked genes in males. Recent studies support this hypothesis. For genes encoding large complexes of seven members or more, which are theoretically dose-sensitive, the expression levels of X-linked genes are equal to those of autosome-linked genes (Pessia et al., 2012). Additionally, it was reported that gene dosage compensation is facilitated by a lower level of m6A modification in X-derived transcripts compared with those linked to autosomes (Rücklé et al., 2023).

Although the term ’gene dosage compensation’ is generally used to refer to a phenomena observed in sex chromosomes, this mechanism is also known to operate in autosomes. Studies have focused on individual genes, such as alcohol dehydrogenase (Birchler, 1979; Birchler, 1981; Birchler and Newton, 1981; Birchler et al., 1990; Guo and Birchler, 1994). Recent advances in transcriptome analyses have enabled the study of gene dosage compensation in organisms with aneuploidy and chromosomal duplications (Graison et al., 2007; Makarevitch and Harris, 2010; Zhang et al., 2010; Hose et al., 2015; Zhang et al., 2017; Krasovec et al., 2023). Studies of autosomal gene dosage compensation in plants have primarily focused on maize (*Zea mays*). Given that both monosomy and trisomy can be produced for all chromosomes in maize, this species is considered tolerant to aneuploidy (Carlson, 1988), facilitating the study of autosomal gene dosage compensation using segmental aneuploidy lines, such as segmental duplications or heterozygous deletions (Yang et al., 2021; Makarevitch et al., 2008). In the case of segmental duplication, analysis of a mutant line, in which 90% of chromosome 5 is trisomic and part of chromosome 6 is monosomic, revealed that gene dosage compensation occurred in some genes within the trisomic region (Makarevitch et al., 2008). The analysis of 17 lines, collectively covering about 80% of all chromosomal regions as disomic, revealed that the imbalance of gene copy numbers are compensated to some degree in expression level (Yang et al., 2021). Similar studies have been performed on the model plant *Arabidopsis thaliana*. The expression analysis of individual trisomic lines for each of the five chromosomes indicated that some genes underwent dosage compensation (Hou et al., 2018). However, in the case of seven segmentally duplicated lines and six heterozygous deletion lines, gene dosage compensation was rarely observed (Picart-Picolo et al., 2020; Tanaka et al., 2020).

If dosage compensation does not occur in heterozygous deletions, the effects of such deletions can influence the phenotype. In the budding yeast *Saccharomyces cerevisiae*, dominant phenotypes have been observed in 40% of the heterozygous loss-of-function mutants of essential genes; this phenomenon is called haploinsufficiency (Ohnuki and Ohya, 2018). A similar phenomena can lead to phenotypic changes in plants. In the present study, we examined the occurrence of gene dosage compensation and its effects on phenotypic changes in 12 *A. thaliana* BC_1_ lines. These lines were produced by crossing mutants with deletions induced by the heavy ion irradiation of wild-type Col-0 plants. Theoretically, heavy-ion beams have the potential to induce large deletions in plants, including *Arabidopsis* (Kazama et al., 2017; Ishii et al., 2024). Thus, evaluating heterozygous deletions is important when considering their usefulness as genetic resources. This study provides a deeper understanding of gene-dosage compensation and phenotypic changes in plants with a heterozygous deletion.

## Materials and Methods

### Plant materials

*A. thaliana* ecotype Columbia (Col-0) and its mutants produced by heavy ion beam irradiation were used in this study (Tables S1 and S2). These plants were grown at 22.5°C under long-day conditions (16 h light/8 h dark) in a growth chamber. Nine of the 12 mutants were previously sequenced, and deletions were detected (Ishii et al., 2024). An additional three mutants, Ar-50-pg1, C200-56-as4, and C30-144-as3, were produced by Ar-ion beam treatment with a linear energy transfer (LET) of 50 keV μm^-1^ at a dose of 50 Gy, C-ion beam treatment with an LET of 200 keV μm^-1^ at a dose of 75 Gy, and C-ion beam treatment with an LET of 30 keV μm^-1^ at a dose of 400 Gy, respectively. To identify these mutants and their deletions, DNA was extracted from fresh leaves using the NucleoSpin Plant II Mini kit for DNA from plants (Macherey-Nagel, Duren, Germany) and sequenced using the HiSeq 4000 sequencing system (Illumina Inc., https://www.illumina.com) as described previously (Kazama et al., 2017). The obtained sequence reads were inputted into the mutation detection pipeline AMAP, as described previously, to detect large deletions (Ishii et al., 2024). From the randomly selected M_2_ plants sequenced, three mutants (Ar-50-pg1, C200-56-as4, and C30-144-as3) were identified to have large deletions. The 12 mutants were crossed with Col-0 plants to produce BC_1_ plants with heterozygous mutations. After crossing, siblings harbouring heterozygous large deletions were selected through genomic PCR on leaves harvested 10 d after cultivation began on 1/2 MS plates containing 1% sugar and 0.7% agar (Murashige and Skoog, 1962). The primers were designed to detect mutant-specific deletions (Table S3). The resulting BC_1_ mutants, each with heterozygous deletions, were used in subsequent analyses.

### Morphological characterization

The selected BC_1_ mutants were grown with the same ageing Col-0 plants one by one in each pot at 22.5°C under long-day conditions (16 h light/8 h dark) in a growth chamber. The flowering times of BC_1_ mutants were measured by counting the number of days until bolting. These morphologies were observed 21 d after the start of cultivation. Each leaf was cut and photographed, and fresh weights were measured.

### RNA-seq analysis

Total RNA was extracted from the leaves of the BC_1_ mutants and Col-0 plants 14 and 40 d after cultivation, and from their flower buds 40 d after cultivation using the Nucleospin Plant and Fungi, Mini kit for RNA from plants and fungi (Macherey-Nagel, Düren, Germany). Strand-specific 3-prime mRNA libraries were prepared using BrAD-seq (Townsley et al., 2015). Briefly, mRNA was isolated from the total RNA using Oligo-d(T)25 Magnetic Beads (New England Biolabs, Ipswich, MA, USA), fragmented by heat and magnesium, and primed for cDNA synthesis using an adapter-containing oligonucleotide. First-strand cDNA was synthesised using RevertAid (Thermo Fisher Scientific, Waltham, MA, USA). The single-stranded portion of a 5-prime adapter was inserted into the breeding terminus of the RNA-cDNA hybrids and incorporated into a complete library molecule using DNA polymerase I (*E. coli*; Takara Bio, Shiga, Japan). PCR enrichment was performed using KOD One PCR Master Mix (TOYOBO, Tokyo, Japan). Libraries were sequenced using an Illumina NovaSeq X Plus 10 B flow cell (Rhelixa Inc., Tokyo, Japan). Only forward reads were mapped to the TAIR10 genome assembly using the STAR aligner with options “--quantMode GeneCounts --outFilterMultimapNmax 1” (version 2.7.10a; Dobin et al., 2013), as reverse reads were of low quality due to the poly A sequences. The differentially expressed genes (DEGs) were called glmQLFit in the R package edgeR (version 3.40.2; Robinson et al., 2010).

## Results

### Development of the heterozygous deletion mutants

To investigate the effects of heterozygous deletions on plant morphology and gene expression in *A. thaliana*, BC_1_ mutants were produced by crossing Col-0 plants and their heavy-ion induced mutants, the mutations of which were detected by whole-genome resequencing, followed by a mutation detection pipeline, AMAP (Ishii et al., 2016; see Materials and Methods). Each BC_1_ mutant was selected by PCR, using primers designed to detect mutant-specific deletions (Table S3). The sizes of the deletions varied from 42.3 Kbp to 2.03 Mbp (Table S2). Genes included in the deletions were identified according to the *Arabidopsis* Information Resource (TAIR: https://www.arabidopsis.org/). Among the BC_1_ mutants, C200-56-as4, Ar-443-as1(1), and C30-144-as3 shared 34 deletions. The 34, 69, and 34 deletions were shared between C200-56-as4 and Ar-443-as1(1), C200-56-as4 and C30-144-as3, and Ar-443-as1(1) and C30-144-as3, respectively (Table S2). In total, heterozygous deletions encompassed 496 genes, with 148 genes commonly deleted in two or more BC1 mutants. Consequently, we investigated the effects of heterozygous deletions on 382 genes (Tables S4–15).

### Effect of heterozygous deletions on plant morphology

To investigate the effects of heterozygous deletions on plant morphology, we examined leaf shape and fresh weight 21 days after the start of cultivation and flowering time in each BC_1_ mutant. Leaf size and shape differed between the BC_1_ mutants and Col-0 plants (Figure 1). Fresh weights also varied between BC_1_ mutants and Col-0 plants (Figure 2a and Table 1); Ar-11-N1, Ar-47-N1, Ar-50-pg1, Ar-94-as1, Ar-443-as1, C100-23-N2, C200-144-as3, and C30-144-as3 showed significantly different fresh weights from Col-0 plants (Figure 2a and Table 1, *p* < 0.05, Student’s *t*-test). Specifically, Ar-11-N1, Ar-50-pg1, Ar-94-as1, C100-23-N2, and C200-144-as3 plants weighed more than Col-0 plants, indicating that heterozygous deletions can have beneficial effects on growth in some cases. These effects were independent of the number of deleted genes (Figure 2; R^2^ = 0.0524).

**Figure 1.**
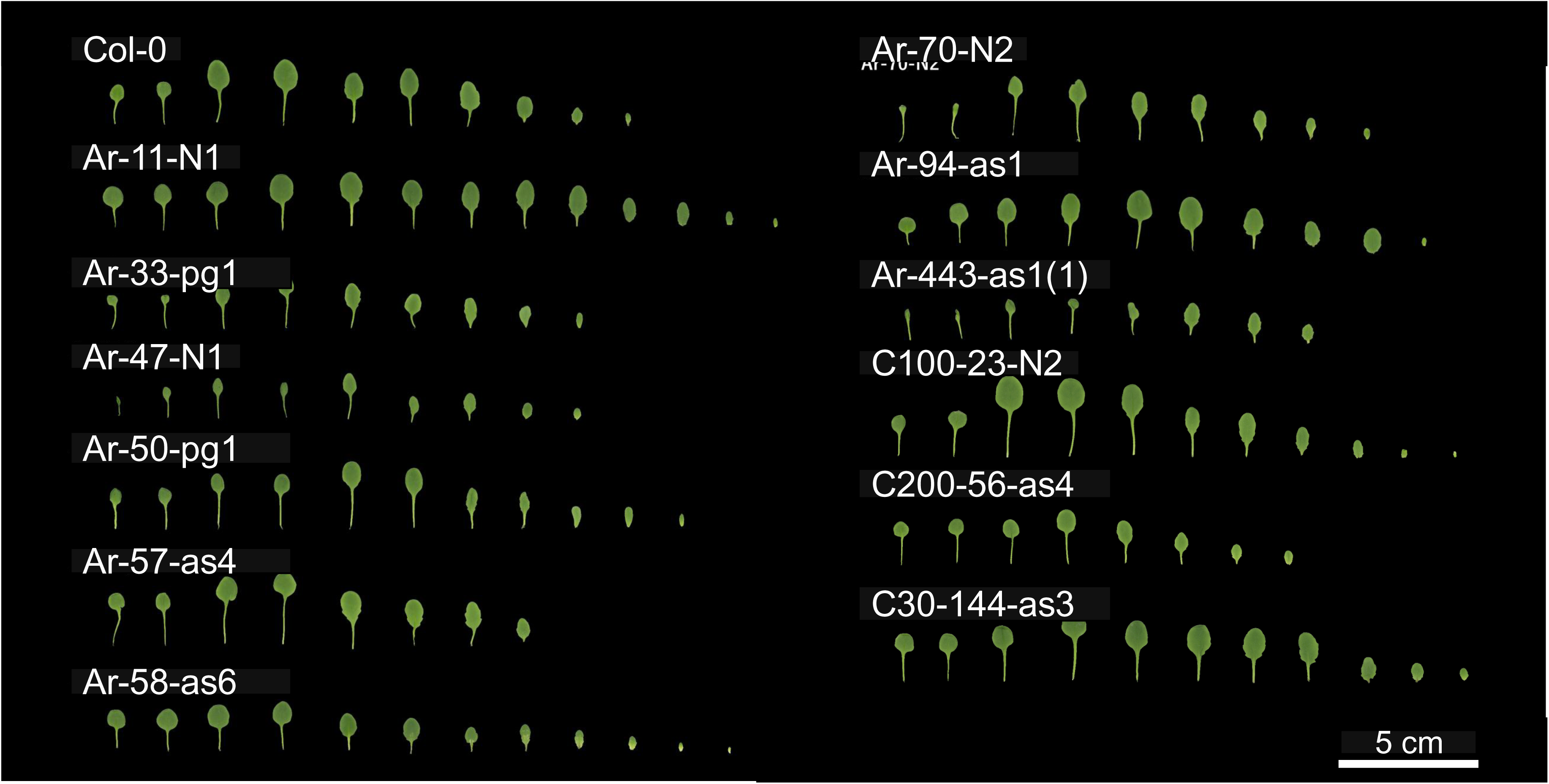
Leaf morphologies of each BC_1_ line on day 21 after cultivation started. Bar = 5 cm.

**Table 1.**
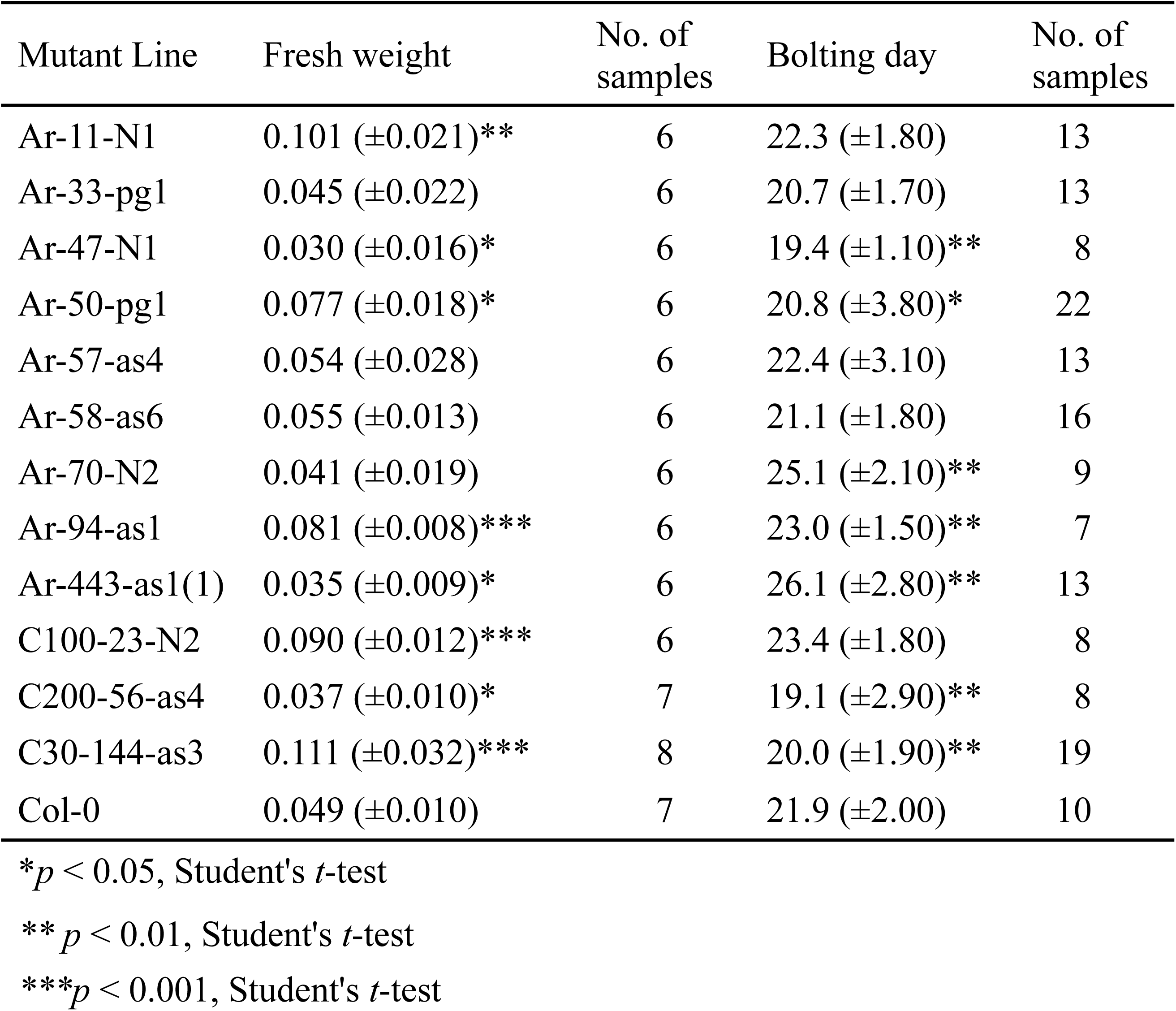
Fresh weight and bolting day in each mutant.

| Mutant Line | Fresh weight | No. of samples | Bolting day | No. of samples |
| --- | --- | --- | --- | --- |
| Ar-11-N1 | 0.101 ($\pm 0.021$)** | 6 | 22.3 ($\pm 1.80$) | 13 |
| Ar-33-pg1 | 0.045 ($\pm 0.022$) | 6 | 20.7 ($\pm 1.70$) | 13 |
| Ar-47-N1 | 0.030 ($\pm 0.016$)* | 6 | 19.4 ($\pm 1.10$)** | 8 |
| Ar-50-pg1 | 0.077 ($\pm 0.018$)* | 6 | 20.8 ($\pm 3.80$)* | 22 |
| Ar-57-as4 | 0.054 ($\pm 0.028$) | 6 | 22.4 ($\pm 3.10$) | 13 |
| Ar-58-as6 | 0.055 ($\pm 0.013$) | 6 | 21.1 ($\pm 1.80$) | 16 |
| Ar-70-N2 | 0.041 ($\pm 0.019$) | 6 | 25.1 ($\pm 2.10$)** | 9 |
| Ar-94-as1 | 0.081 ($\pm 0.008$)*** | 6 | 23.0 ($\pm 1.50$)** | 7 |
| Ar-443-as1(1) | 0.035 ($\pm 0.009$)* | 6 | 26.1 ($\pm 2.80$)** | 13 |
| C100-23-N2 | 0.090 ($\pm 0.012$)*** | 6 | 23.4 ($\pm 1.80$) | 8 |
| C200-56-as4 | 0.037 ($\pm 0.010$)* | 7 | 19.1 ($\pm 2.90$)** | 8 |
| C30-144-as3 | 0.111 ($\pm 0.032$)*** | 8 | 20.0 ($\pm 1.90$)** | 19 |
| Col-0 | 0.049 ($\pm 0.010$) | 7 | 21.9 ($\pm 2.00$) | 10 |

\* $p < 0.05$ , Student's $t$ -test
\*\* $p < 0.01$ , Student's $t$ -test
\*\*\* $p < 0.001$ , Student's $t$ -test

**Figure 2.**
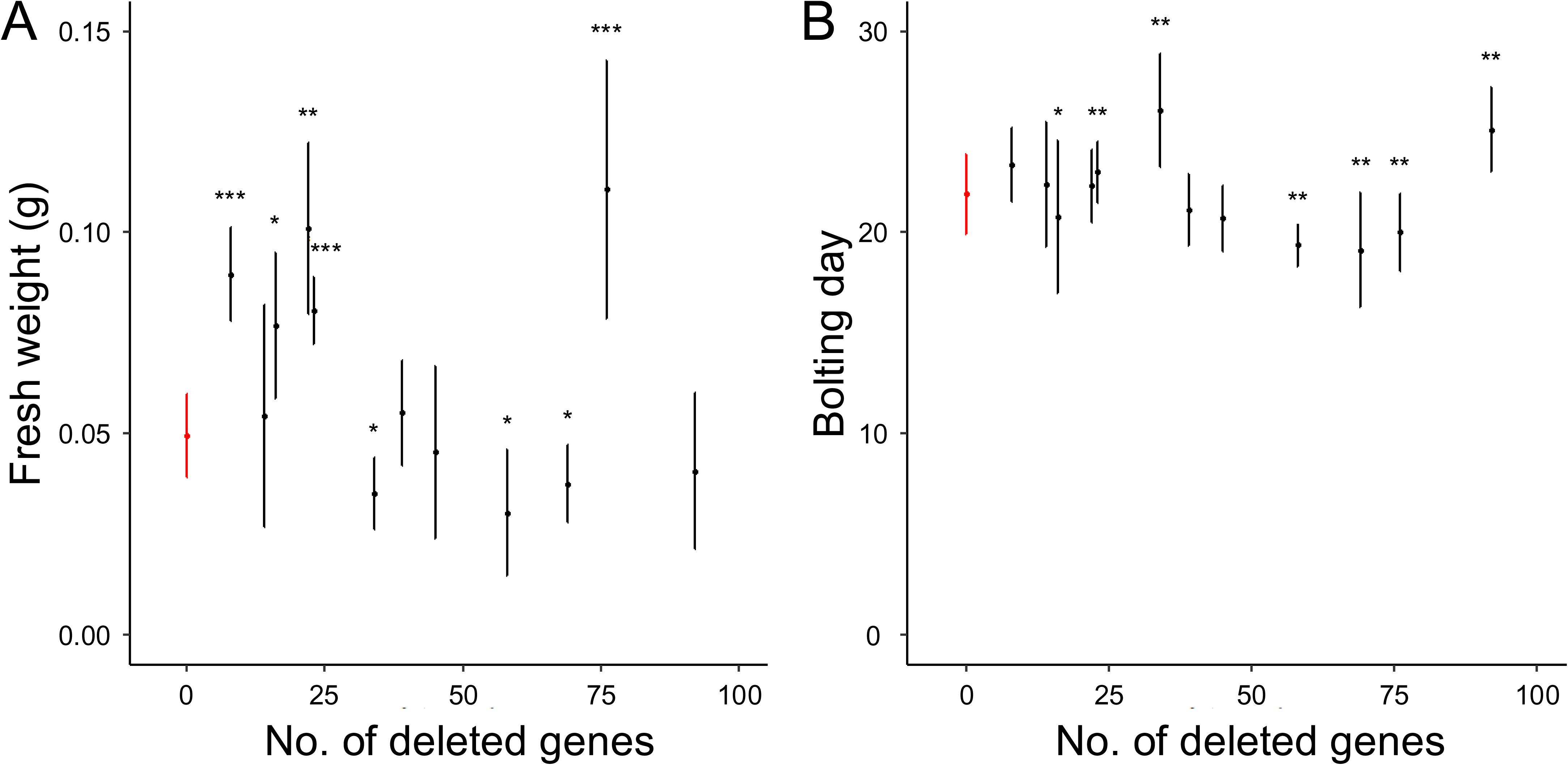
Relation between the number of deleted genes and fresh weights (A) and bolting days (B). Red and black plots represent Col-0 and mutants, respectively. *, *p* < 0.05; **, *p* < 0.01; and ***, *p* < 0.001 (Student’s *t*-test).

Flowering times also varied between BC_1_ mutants and Col-0 plants (Figure 2b and Table S1); Ar-47-N1, Ar-50-pg1, C200-56-as4, and C30-144-as3 bolted faster than Col-0 plants (*p* < 0.05, Wilcoxon rank-sum test), whereas Ar-70-N2, Ar-94-as1, and Ar-443-as1 bolted slower than Col-0 plants (*p* < 0.05, Wilcoxon rank-sum test). Flowering times were also independent of the number of deleted genes (Figure 2, R^2^ = 0.0273). Collectively, nine of the 12 BC_1_ mutants tested showed morphological changes, indicating that heterozygous deletions can cause phenotypic changes in *A. thaliana*.

### Effect of heterozygous deletions on gene expression

To investigate the effect of heterozygous deletions on gene expression, mRNA was extracted from three independent plants each of the BC_1_ mutant and Col-0, from whole plants 14 d after cultivation started, and from leaves and flower buds 40 d after cultivation started. RNA-seq analysis was performed using a NextSeq 500 (see Materials and Methods). When the gene expression ratios of the BC_1_ mutant to Col-0 for each gene in the non-deleted regions were calculated, including all mutant samples, the peak of their density was observed at approximately one, indicating no difference in gene expression levels between the BC_1_ mutants and Col-0 plants (Figure 3a). By contrast, when the ratios were calculated for genes in the heterozygous deletion regions of all mutant samples, the peak density was observed at around 0.5, indicating that the expression levels of genes in heterozygous deletions were halved (Figure 3b). The dispersion of densities was significantly different between the non-deleted and deleted regions (*p* < 0.001, Kolmogorov–Smirnov test). Therefore, gene dosage compensation was not observed for most of the genes with heterozygous deletions tested.

**Figure 3.**
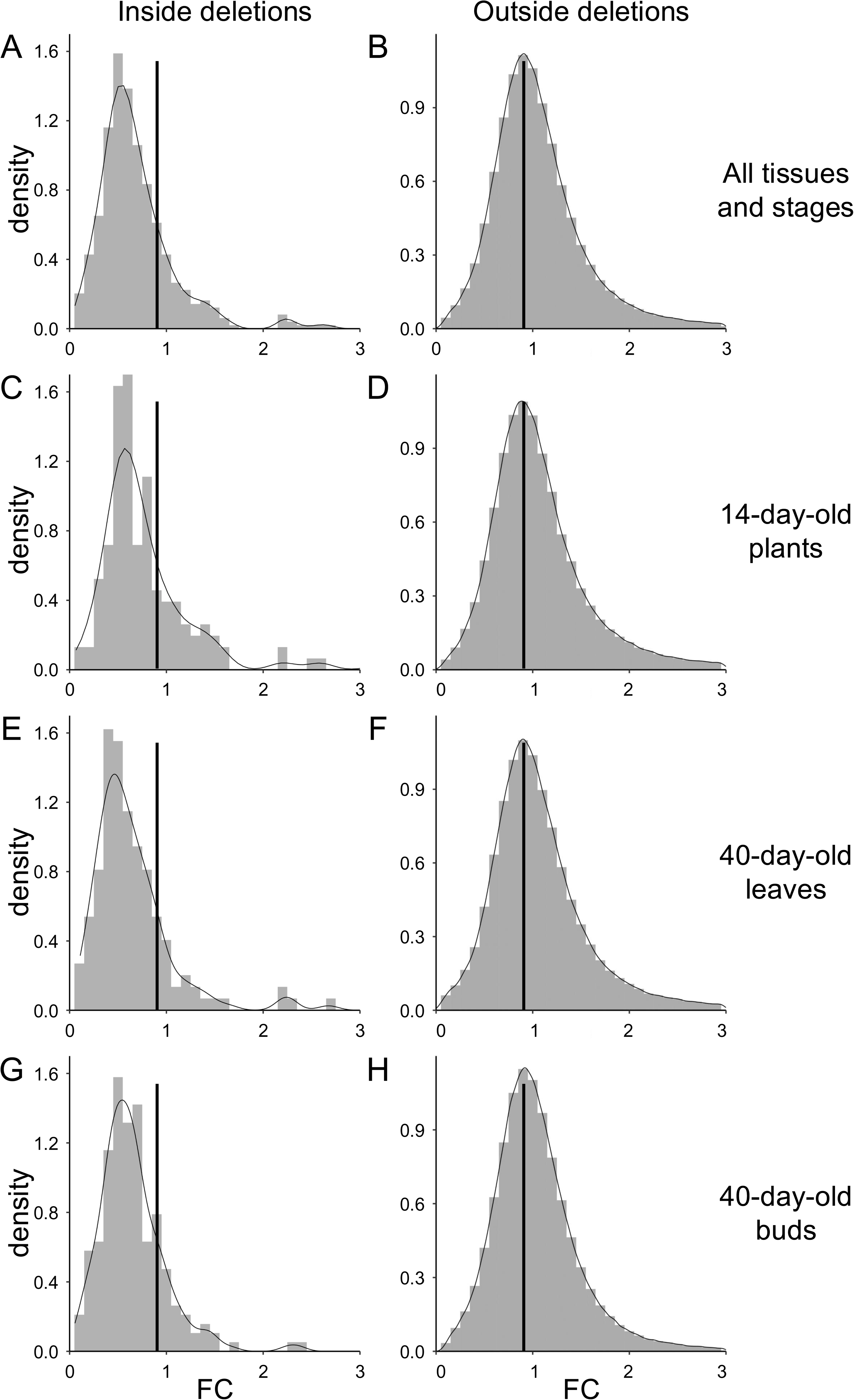
Histograms of gene expression ratios of BC_1_ mutant to Col-0 for genes located outside (A, C, E, G) and located inside the deletions (B, D, F, H). A and B: Histograms depicting all samples. C and D: Histograms depicting plants on day 14 after cultivation started. E and F: Histograms depicting leaves on day 40 after cultivation started. G and H: Histograms depicting flower buds on day 40 after cultivation started.

The expression levels of genes located within the heterozygous deletion regions in each BC_1_ mutant were investigated and compared with the expression levels of the same genes in Col-0 for each tissue and developmental stage. The results indicated that the median expression levels were significantly reduced in BC_1_ mutants across all tissues and stages (Figure S1, *p* < 0.05, Welch’s t-test), except for whole plants at 14 d post-cultivation, where the difference was not statistically significant (Figure S1, *p* > 0.05, Welch’s *t*-test); although the mutants exhibited lower expression levels. The histogram of the gene expression ratios within the deletion regions of the BC_1_ mutant to Col-0 for each gene also showed peaks at approximately 0.5 (Figure 3c–h), whereas those outside the region showed peaks at approximately one (*p* < 0.001, Kolmogorov–Smirnov test). These results indicated that gene dosage compensation did not occur in most genes at different developmental stages. However, the histograms showed small peaks around 0.8 in whole plants 14-d after cultivation started and flower buds 40 d after cultivation started (Figure 3d and h), suggesting the possibility of gene dosage compensation in a small number of genes. The genes undergoing dosage compensation were not strain-or deletion-specific but were scattered across various strains or deletions. Therefore, no deletion specificity is observed in the presence of dosage compensation. When the data were presented in MA plots, increases in expression ratios were observed regardless of the average expression level in all tissues and stages (Figure S2). Thus, the increase in expression levels in the mutants was not due to the variability in data caused by genes with low expression levels.

When the copy number of a specific gene is altered, the equilibrium of expression levels between the gene and its partner genes (such as genes encoding subunits of the same complex) is disrupted, resulting in adverse effects on cellular function. These genes are known as Dosage Balance Genes (DBGs; Papp et al., 2003). The DBGs tend to be maintained with their partner genes at dosage balances after whole-genome duplication. Therefore, paralogs that underwent whole-genome duplication were regarded as DBGs in *A. thaliana* (Bowers et al., 2003; Thomas et al., 2006; Tables S16–22). We examined whether dosage compensation occurred in the DBGs included in the heterozygous BC1 deletion mutants. In total, 50 DBGs were included in the heterozygous deletions. A comparison of the expression levels of DBGs in the heterozygous deletions between Col-0 and BC_1_ mutant plants revealed that the median expression levels were significantly lower in the BC_1_ mutants (Figure 4, *p* < 0.05, Welch’s *t*-test). This trend persisted when the median expression levels were examined across three distinct developmental stages. When the expression levels of non-deleted paralogs of heterozygously deleted DBGs were compared between Col-0 plants and BC1 mutants, no significant differences were observed in the median expression levels (Figure S3, *p* > 0.05, Welch’s *t*-test). This result indicates that the upregulation of DBGs does not occur when the copy numbers of their paralog genes are halved.

**Figure 4.**
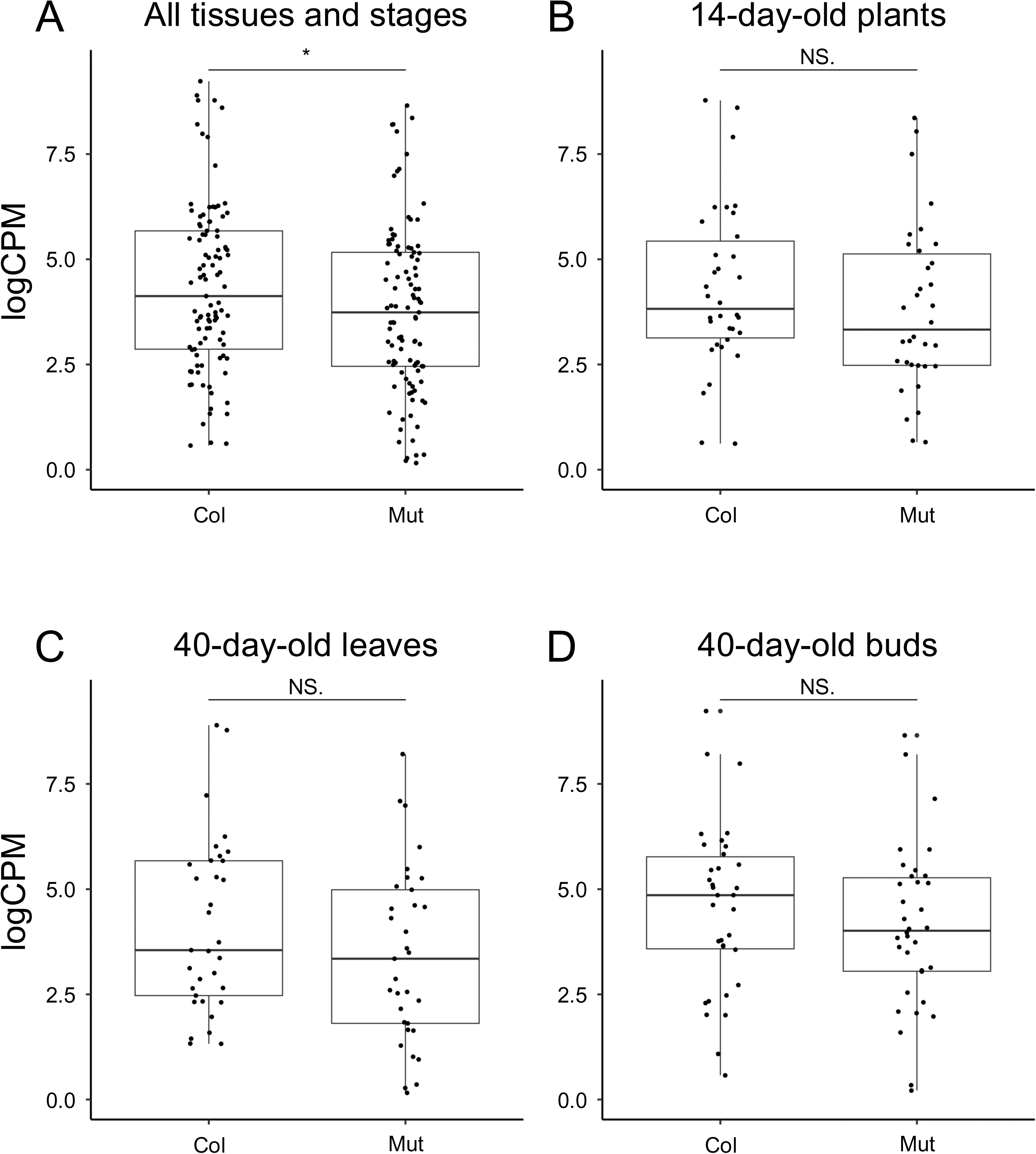
Box-plot comparing Col-0 and the mutant for the expression level of the Dosage Balance Genes (DBGs) inside the deletions in all samples tested (A), in plants at day 14 after cultivation started (B), in leaves on day 40 after cultivation started (C), and in buds on day 40 after cultivation started (D). NS; *p* > 0.05 (Welch’s t-test).

Ar-443-as1(1), C30-144-as3, and C200-56-as1 exhibited heterozygous gene deletions in the same chromosomal region, which facilitated the detection of consistent upregulation of these genes (Table S23). Notably, AT3G29639 and AT3G29770 showed elevated expression at all sampling stages for both C200-56-as4 and C30-144-as3. AT3G30160 and AT3G30460 showed increased expression exclusively in flower buds and young leaves (14 d after cultivation), respectively. None of these genes encode transcription factors or proteins known to form specific complexes, and their functions remain unidentified. Therefore, a small number of genes may be dose-compensated in a gene-by-gene manner.

## Discussion

Compared to whole-genome gene duplication, heterozygous deletions or segmental duplications may influence the balance of gene expression in the genome, as in the case of X-or Z-linked genes on sex chromosomes. This imbalance can cause phenotypic changes in the absence of gene dosage compensation. In the present study, the effects of heterozygous deletions on phenotypic changes and gene expression levels were investigated using 12 BC_1_ *A. thaliana* mutants produced by crossing Col-0 plants with heavy ion-induced mutants with large deletions.

In *A. thaliana,* dosage compensation does not occur in cases of trisomies, nor in a trans effect in which the expression levels of non-trisomic chromosomes change (Hou et al., 2018). BC_1_ mutants tested in the current study did not show gene dosage compensation, corresponding to a previous report in which six different mutants produced by the ExTaqing system were tested (Tanaka et al., 2020). Here, we also demonstrated that dosage compensation did not occur in most of the heterozygously deleted genes across the three developmental stages (Figure 3). Additionally, our data showed that dosage compensation did not occur in these genes, regardless of their expression levels (Figure S2). These observations contrast with those in *Drosophila*, wherein gene dosage compensation was observed in genes with high expression ratios in proportion to the average gene expression levels when heterozygous deletions occurred (McAnally and Yampolsky, 2010). In the case of heterozygous deletions, we also observed that the trans-effect did not occur (Figure 3). The absence of a trans-effect may result from a lack of transcription factors. In our dataset, 22 of 496 genes encoded transcription factors, and no trans effect was observed in any of the BC_1_ mutants (Tables S4–15). Deletion size is also a possible cause of the absence of trans effects because the number of hemizygous genes is restricted by the deletion size. Indeed, in the case of disomic haploid maize lines where both trans effect and compensation were observed in some extent, in which 876 or more deleted genes were included in individual disomic regions (Yang et al., 2021). The number of hemizygous genes is restricted by the deletion sizes. However, the size of deletions induced by heavy-ion irradiation is restricted by the distribution of essential genes in the haploid phase (Ishii et al., 2024). Taken together, we conclude that gene dosage compensation does not occur in most genes when heterozygous deletions are induced by heavy-ion irradiation in *A. thaliana*.

Gene dosage compensation has been observed in the heteromorphic sex chromosomes of six plant species: *Silene latifolia*, *Cannabis sativa*, *Humulus lupulus*, *Coccinia grandis*, *Rumex hastatulus*, and *Rumex rothschildianus* (Hough et al., 2014; Papadopulous et al., 2015; Crowson et al., 2017; Fruchard et al., 2020; Prentout et al., 2020; Prentout et al., 2021; Muyle et al., 2022). This phenomenon likely balances the expression of dosage-sensitive genes, thereby compensating for the degeneration of Y- or W-linked genes. In *S. latifolia*, heavy-ion irradiation-induced large deletions in the Y chromosome led to the upregulation of gene expression levels in X-linked genes homologous to the deleted Y-linked genes, a strict process termed immediate dosage compensation (Krasovec et al., 2019). To explore the evolution of gene dosage compensation in plants, replicating the same experiment on autosomes could serve as a counterpart for the *S. latifolia* study. Our results indicated that immediate dosage compensation does not occur in *A. thaliana* autosomes (Figure 3). Nonetheless, we found evidence of dosage compensation in some genes (Table S23), implying that these potentially dosage-sensitive genes may contribute to the development of gene-by-gene dosage compensation mechanisms in plant sex chromosomes.

Our current findings indicate that nine of the 12 BC_1_ mutants exhibited morphological changes (Figure 1, Table 1). Notably, five BC_1_ mutants demonstrated increased fresh weight 14 d after the start of cultivation. These observations suggest that substantial heterozygous deletions could serve as genetic resources for breeding despite the challenges in identifying the specific genes responsible for the observed phenotypic alterations. Such heterozygous deletions can be efficiently generated using heavy-ion beam irradiation with high LET values (Hirano et al., 2015; Kazama et al., 2017). Heavy-ion beam irradiation is applied to various tissues, organs, and plant species (Abe et al., 2015). Consequently, high-LET heavy-ion irradiation has emerged as a potent tool for inducing heterozygous deletions, leading to phenotypic variations that may differ from those caused by the complete disruption of a single gene.

### Data availability

Data were deposited with links to BioProject numbers PRJDB17829 for whole-genome sequencing and PRJDB17834 for RNA-seq of the mutants in the DDBJ BioProject database (https://ddbj.nig.ac.jp/search/en).

## Acknowledgements

We thank RIKEN Nishina Centre and the Centre for Nuclear Study, University of Tokyo, for the operation of RIBF that enabled us to perform ion-beam irradiation. We would also like to thank Dr. Atsushi Yoshida (RIKEN) for calculating the penetration distances of heavy-ion beams.

## Author contributions

TI, RN, MI, and ADN performed the experiments and obtained the data presented in this manuscript. TI, RN, and KI performed the data analysis. TA conducted the heavy-ion beam irradiation. YK designed the study. TI and YK wrote the first draft of the manuscript. All the authors reviewed the draft and approved the final version of the manuscript.

---

**Figure S1.**
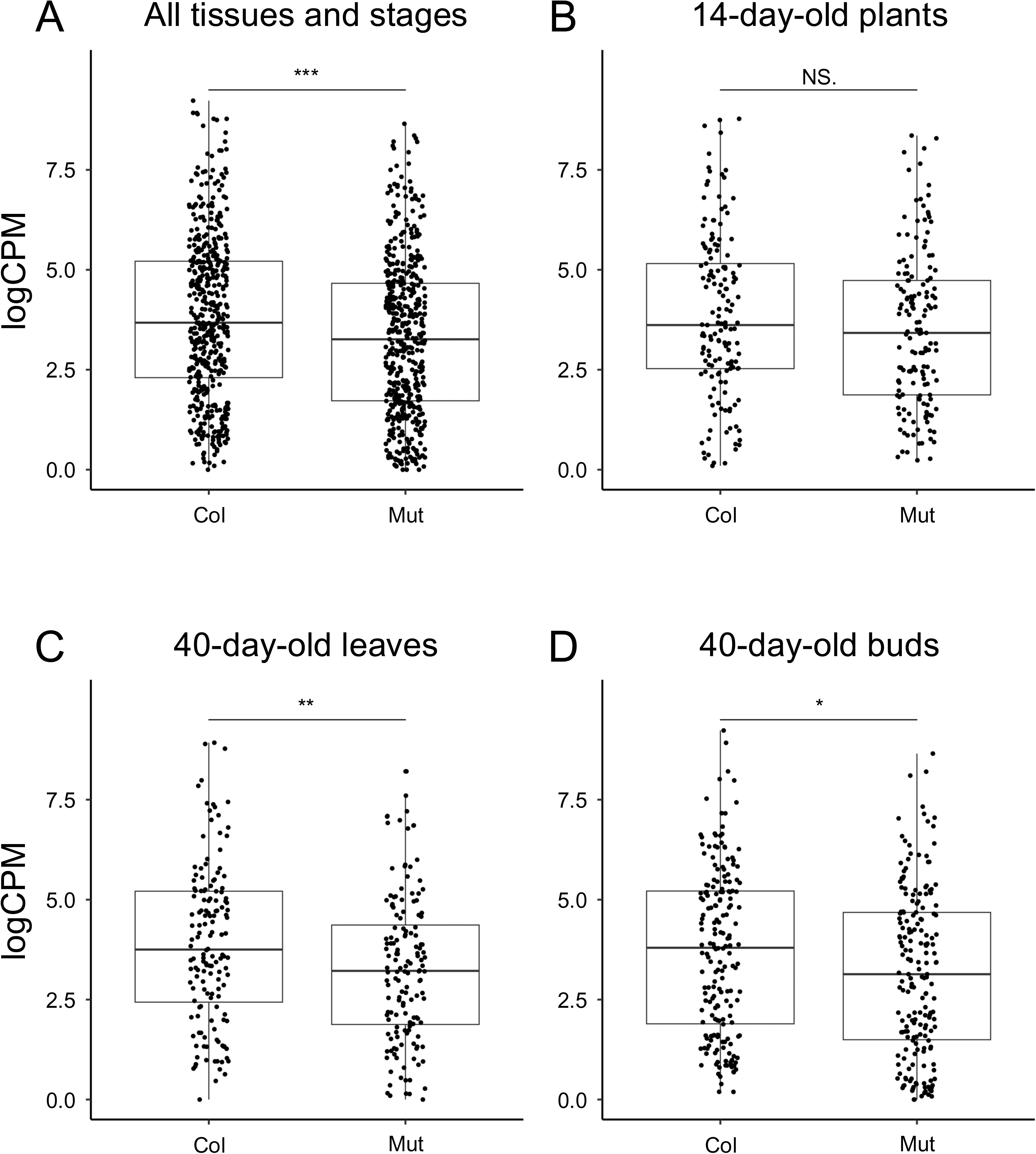
Comparison of gene expression levels inside the deletions between Col-0 and the mutant for all tissues and stages (A), 14-day-old plants (B), 40-day-old leaves (C), and 40-day-old flower buds (D). Statistical significance is indicated as \*\*\**p* < 0.001, \*\**p* < 0.01, \**p* < 0.05, and ns (not significant) (Welch’s t-test).

**Figure S2.**
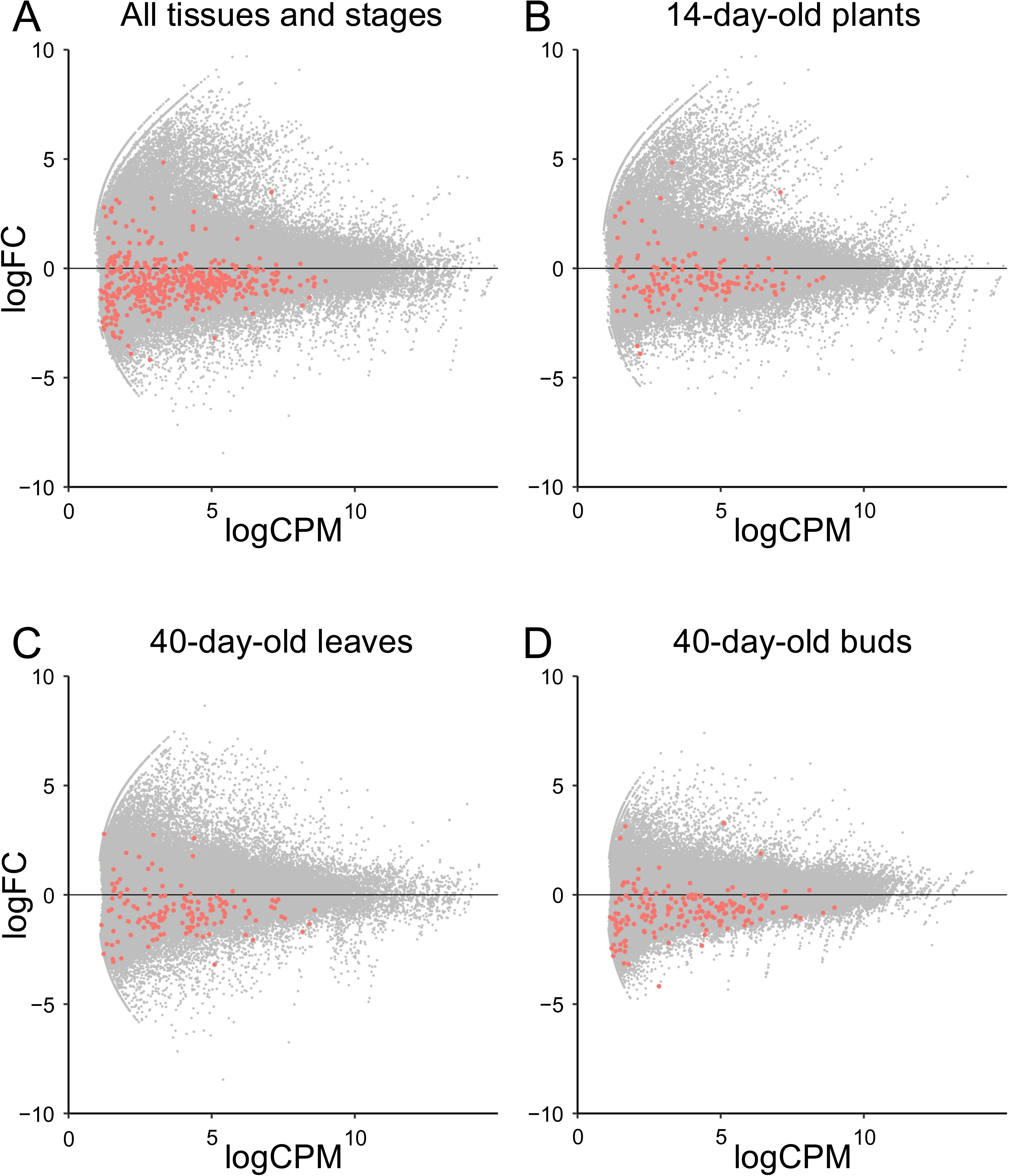
MA plot with mean gene expression on the horizontal axis and gene expression ratio of mutants to Col-0 on the vertical axis, for all tissues and stages (A), 14-day-old plants (B), 40-day-old leaves (C), and 40-day-old flower buds (D). Red plots indicate intra-deletion genes, and grey plots indicate extra-deletion genes.

**Figure S3.**
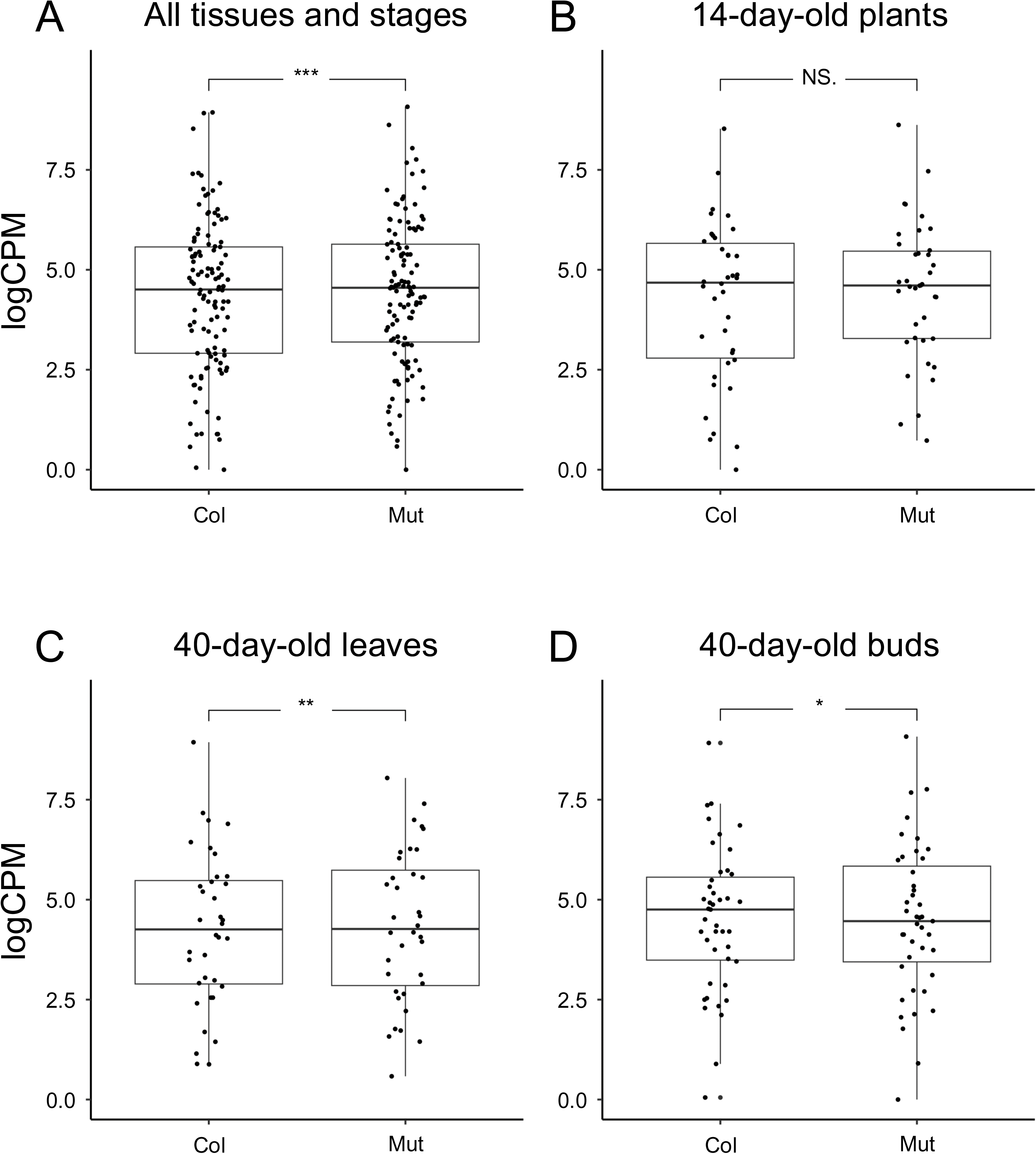
Comparison of expression levels of Dosage Balance Genes (DBGs) present outside the deletion paired with DBGs inside the deletions between Col-0 and the mutant for all tissues and stages (A), 14-day-old plants (B), 40-day-old leaves (C), and 40-day-old flower buds (D). No significant differences were detected (Welch’s t-test).

**Table S1.**
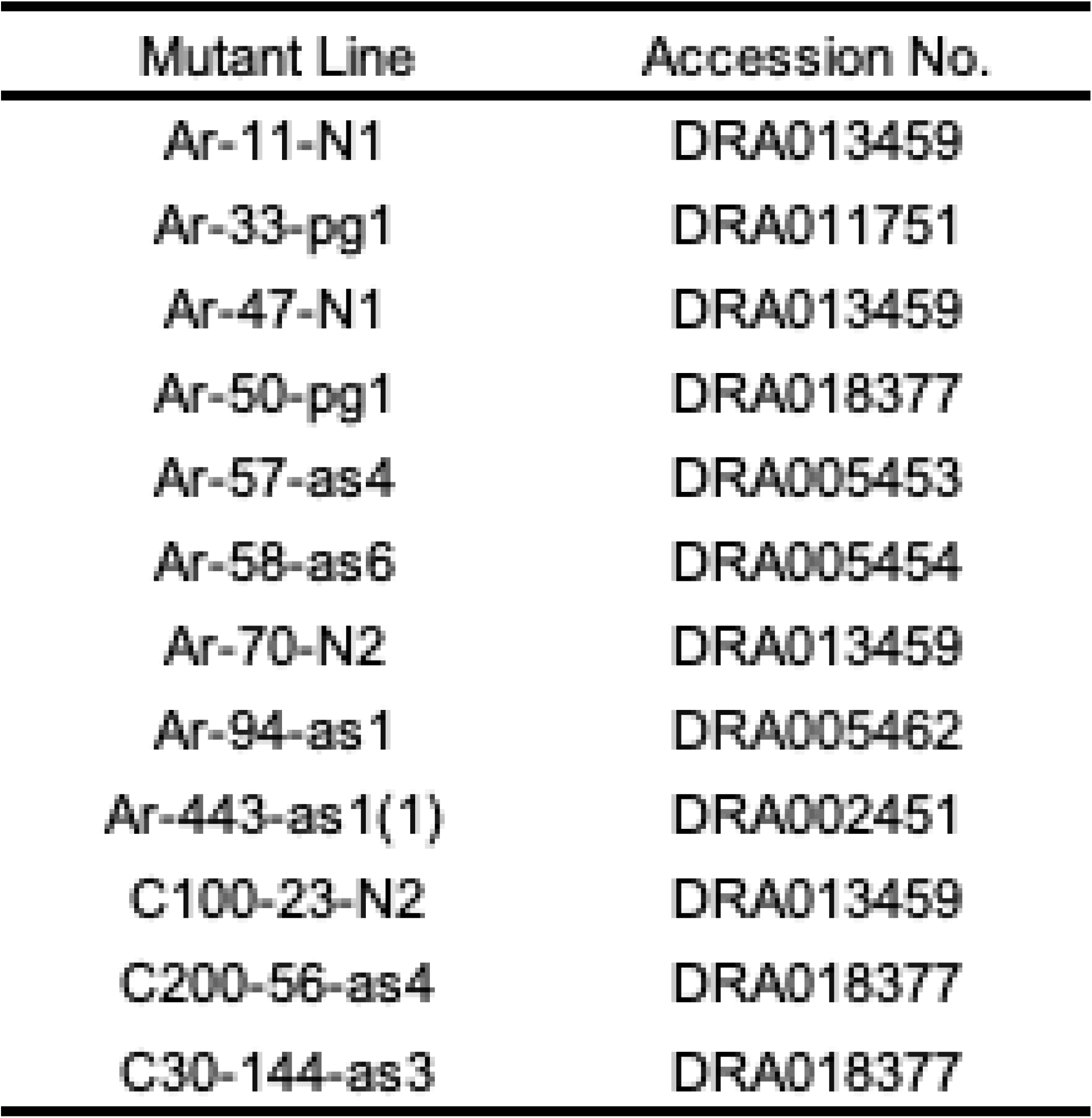
Accession number for each mutant

**Table S2.** Deletion mutations analyzed in this study.

| Mutant Line | Chromo-<br>some | Start position<br>(bp) | End position<br>(bp) | Deletion<br>size (bp) | Number of<br>deleted gene | Reference |
| --- | --- | --- | --- | --- | --- | --- |
| Ar-11-N1 | 5 | 15,036,948 | 15,170,720 | 133,773 | 22 | Ishi et al. 2024 |
| Ar-33-pg1 | 5 | 12,621,533 | 13,561,538 | 940,006 | 45 | Sanjaya et al. 2021 |
| Ar-47-N1 | 3 | 16,980,794 | 17,226,677 | 245,884 | 58 | Ishi et al. 2024 |
| Ar-50-pg1 | 5 | 12,578,194 | 13,032,663 | 454,470 | 16 | This study |
| Ar-57-as4 | 2 | 9,483,679 | 9,534,538 | 50,860 | 14 | Kazama et al. 2017 |
| Ar-58-as6 | 5 | 10,068,563 | 10,544,692 | 476,130 | 39 | Kazama et al. 2017 |
| Ar-70-N2 | 1 | 13,057,211 | 15,084,979 | 2,027,769 | 92 | Ishi et al. 2024 |
| Ar-94-as1 | 1 | 23,422,006 | 23,494,138 | 72,133 | 23 | Ishi et al. 2024 |
| Ar-443-as1(1) | 3 | 11,761,164 | 12,363,310 | 602,147 | 34 | Hirano et al. 2015 |
| C100-23-N2 | 3 | 10,377,094 | 10,419,440 | 42,347 | 8 | Ishi et al. 2024 |
| C200-56-as4 | 3 | 11,390,812 | 12,570,716 | 1,179,905 | 69 | This study |
| C30-144-as3 | 3 | 11,316,869 | 12,608,684 | 1,291,816 | 76 | This study |

**Table S3.**
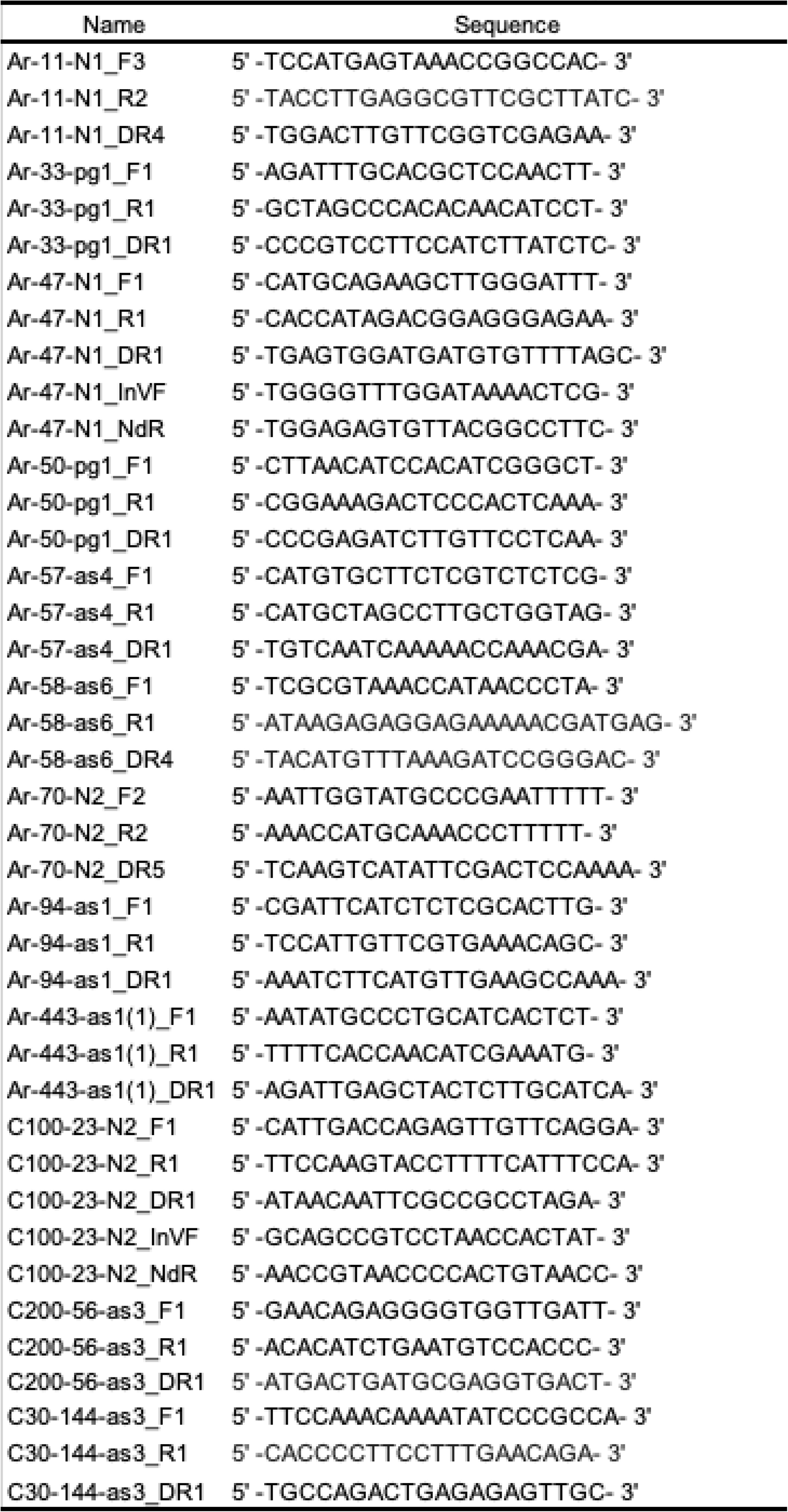
Primers used for deletion detection in this study.

**Table S4.**
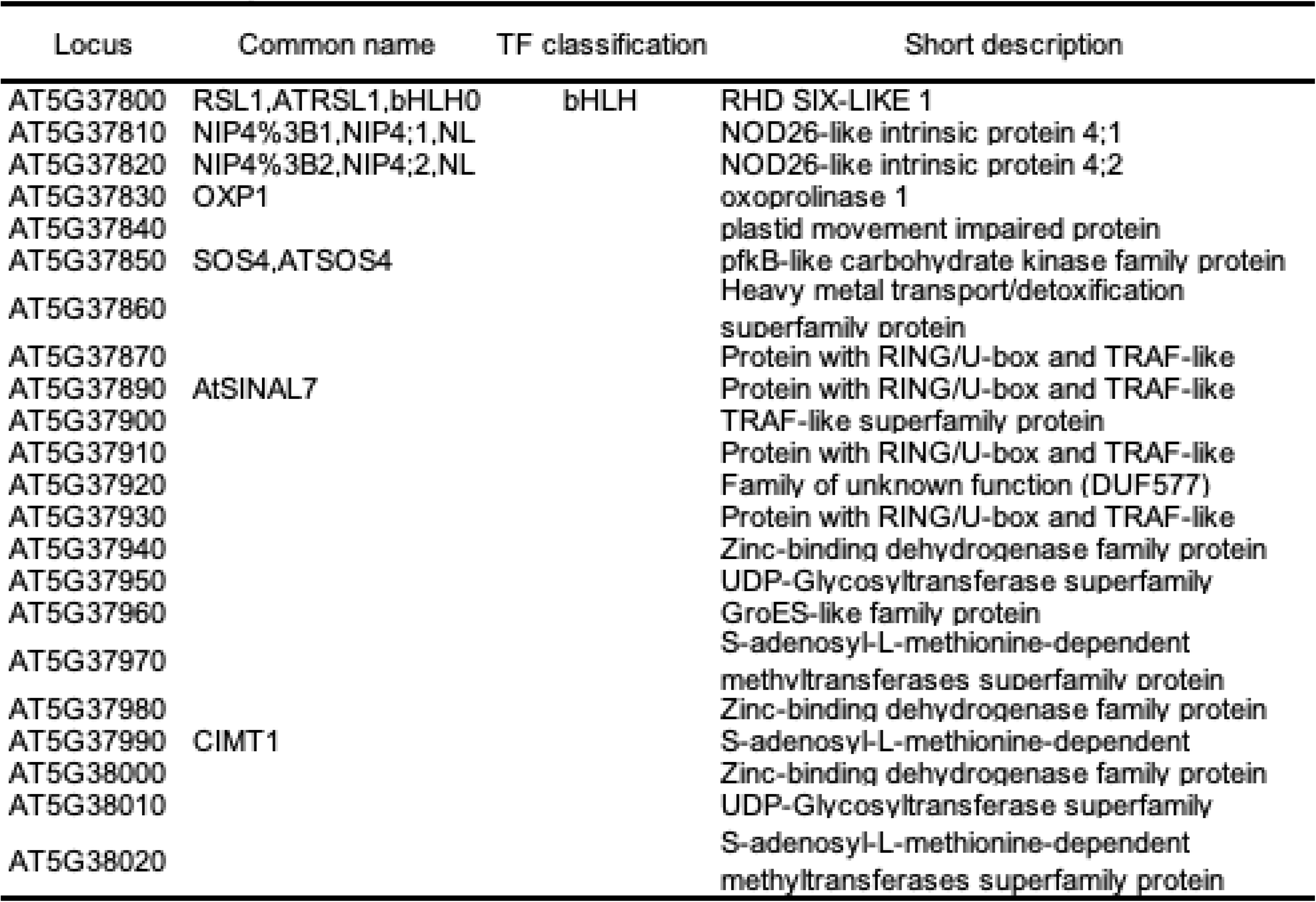
Deleted genes in AM 1-N1.

**Table S5.**
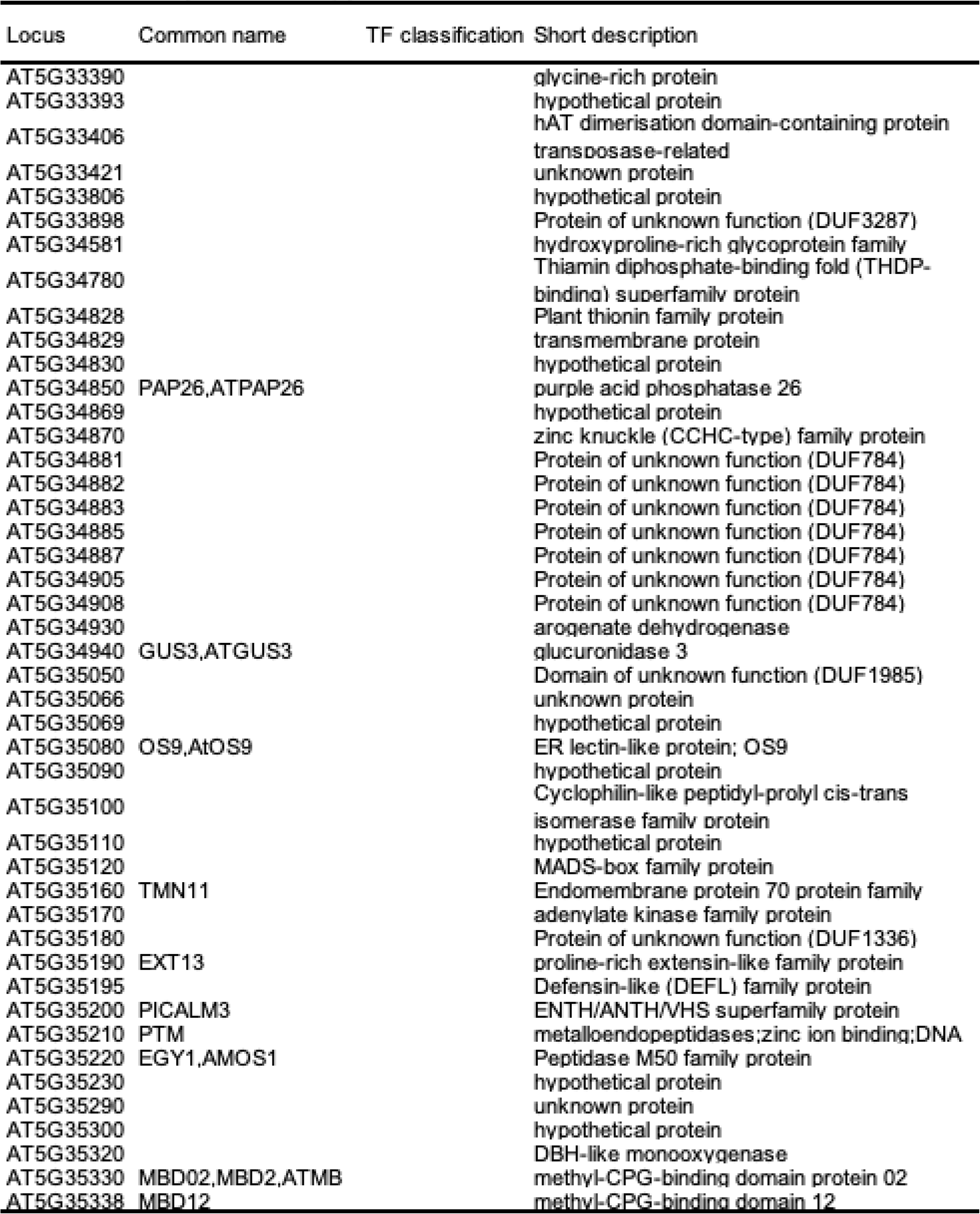
Deleted genes in Ar-33-pg1.

**Table S6.**
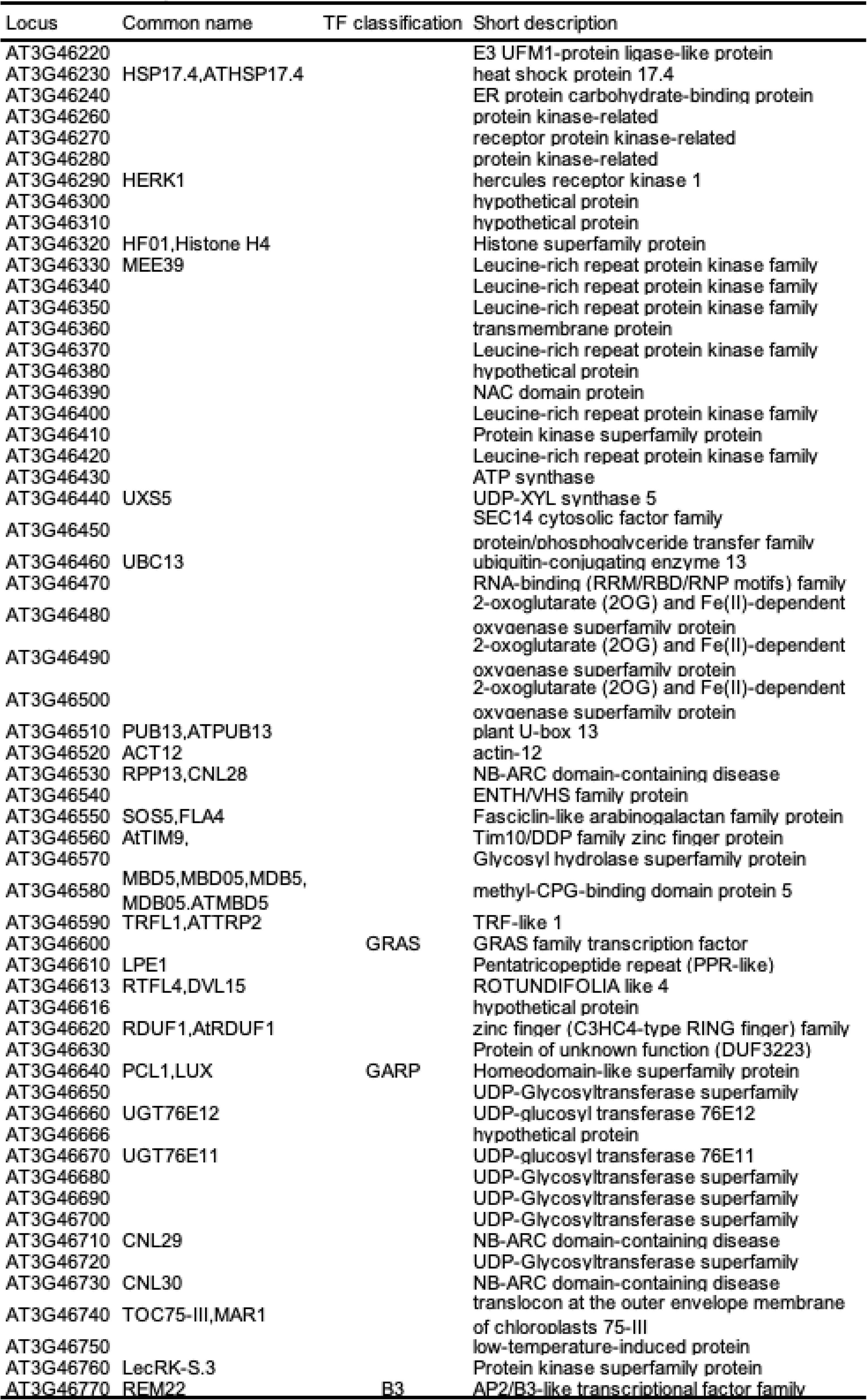
Deleted genes in Ar-47-N1

**Table S7.** Deleted genes in Ar-50-pg 1

| Locus | Common name | TF classification | Short description |
| --- | --- | --- | --- |
| AT5G33320 | NOX1, |  | Glucose-6-phosphate/phosphate |
| AT5G33330 |  |  | Polynucleotidyl transferase, ribonuclease H-like superfamily protein |
| AT5G33340 | CDR1 |  | Eukaryotic aspartyl protease family protein |
| AT5G33355 |  |  | Defensin-like (DEFL) family protein |
| AT5G33370 | CUS2 |  | GDSL-like Lipase/Acylhydrolase superfamily |
| AT5G33390 |  |  | glycine-rich protein |
| AT5G33393 |  |  | hypothetical protein |
| AT5G33406 |  |  | hAT dimerisation domain-containing protein/transposase-related |
| AT5G33421 |  |  | unknown protein |
| AT5G33806 |  |  | hypothetical protein |
| AT5G33898 |  |  | Protein of unknown function (DUF3287) |
| AT5G34581 |  |  | hydroxyproline-rich glycoprotein family |
| AT5G34780 |  |  | Thiamin diphosphate-binding fold (THDP-binding) superfamily protein |
| AT5G34828 |  |  | Plant thionin family protein |
| AT5G34829 |  |  | transmembrane protein |
| AT5G34830 |  |  | hypothetical protein |

**Table S8.** Deleted genes in Ar-57-as4.

| Locus | Common name | TF classification | Short description |
| --- | --- | --- | --- |
| AT2G22330 | CYP79B3 |  | cytochrome P450, family 79, subfamily B, polypeptide 3 |
| AT2G22340 |  |  | transmembrane protein |
| AT2G22345 |  |  | Cysteine-rich protein |
| AT2G22360 | DJA6 |  | DNAJ heat shock family protein |
| AT2G22370 | Med18 |  | mediator of RNA polymerase II transcription |
| AT2G22400 | AtTRM4b |  | S-adenosyl-L-methionine-dependent methyltransferases superfamily protein |
| AT2G22410 | SLO1 |  | SLOW GROWTH 1 |
| AT2G22420 | PRX17 |  | Peroxidase superfamily protein |
| AT2G22425 |  |  | Microsomal signal peptidase 12 kDa subunit |
| AT2G22426 |  |  | hypothetical protein |
| AT2G22430 | HB6,ATHB6 |  | homeobox protein 6 |
| AT2G22440 |  |  | non-LTR retroelement reverse transcriptase |
| AT2G22450 | RIBA2,AtRIBA2 |  | riboflavin biosynthesis protein, putative |
| AT2G22460 |  |  | Protein of unknown function, DUF617 |

**Table S9.** Deleted genes in Ar-58-as6.

| Locus | Common name | TF classification | Short description |
| --- | --- | --- | --- |
| AT5G28060 | RPS24B |  | Ribosomal protein S24e family protein |
| AT5G28070 |  |  | unknown protein |
| AT5G28080 | AtWNK9, WNK9 |  | Protein kinase superfamily protein |
| AT5G28090 |  |  | hypothetical protein |
| AT5G28150 |  |  | Plant protein of unknown function (DUF868) |
| AT5G28160 |  |  | Galactose oxidase/kelch repeat superfamily |
| AT5G28180 |  |  | Galactose oxidase/kelch repeat superfamily |
| AT5G28190 |  |  | transmembrane protein |
| AT5G28210 |  |  | mRNA capping enzyme family protein |
| AT5G28220 |  |  | Protein prenyltransferase superfamily |
| AT5G28235 |  |  | Ulp1 protease family protein |
| AT5G28237 |  |  | Pyridoxal-5'-phosphate-dependent enzyme family protein |
| AT5G28288 |  |  | Defensin-like (DEFL) family protein |
| AT5G28290 | NEK3,ATNEK3 |  | NIMA-related kinase 3 |
| AT5G28295 |  |  | hypothetical protein |
| AT5G28300 | AtGT2L | Trihelix | Duplicated homeodomain-like superfamily |
| AT5G28310 |  |  | NAD(P)-binding Rossmann-fold superfamily |
| AT5G28320 |  |  | embryo defective protein |
| AT5G28340 |  |  | Tetratricopeptide repeat (TPR)-like |
| AT5G28350 |  |  | Quinoprotein amine dehydrogenase, beta chain-like: RIC1-like |
| AT5G28370 |  |  | Pentatricopeptide repeat (PPR) superfamily |
| AT5G28380 |  |  | Tetratricopeptide repeat (TPR)-like |
| AT5G28390 |  |  | RNA-binding (RRM/RBD/RNP motifs) family |
| AT5G28400 |  |  | embryo defective protein |
| AT5G28410 |  |  | hypothetical protein |
| AT5G28420 |  |  | Pentatricopeptide repeat (PPR) superfamily |
| AT5G28442 |  |  | quinoprotein amine dehydrogenase, beta chain-like: RIC1-like |
| AT5G28450 |  |  | Chlorophyll A-B binding family protein |
| AT5G28460 |  |  | Pentatricopeptide repeat (PPR) superfamily |
| AT5G28463 |  |  | transmembrane protein |
| AT5G28465 |  |  | unknown protein |
| AT5G28470 |  |  | Major facilitator superfamily protein |
| AT5G28490 | LSH1,OBO2 |  | Protein of unknown function (DUF640) |
| AT5G28491 |  |  | hypothetical protein |
| AT5G28500 |  |  | rubisco accumulation factor-like protein |
| AT5G28510 | BGLU24 |  | beta glucosidase 24 |
| AT5G28520 |  |  | Mannose-binding lectin superfamily protein |
| AT5G28530 | FRS10 | FAR1 related | FAR1-related sequence 10 |
| AT5G28540 | BIP1,MED37,Med37 |  | heat shock protein 70 (Hsp 70) family protein |

**Table S10.**
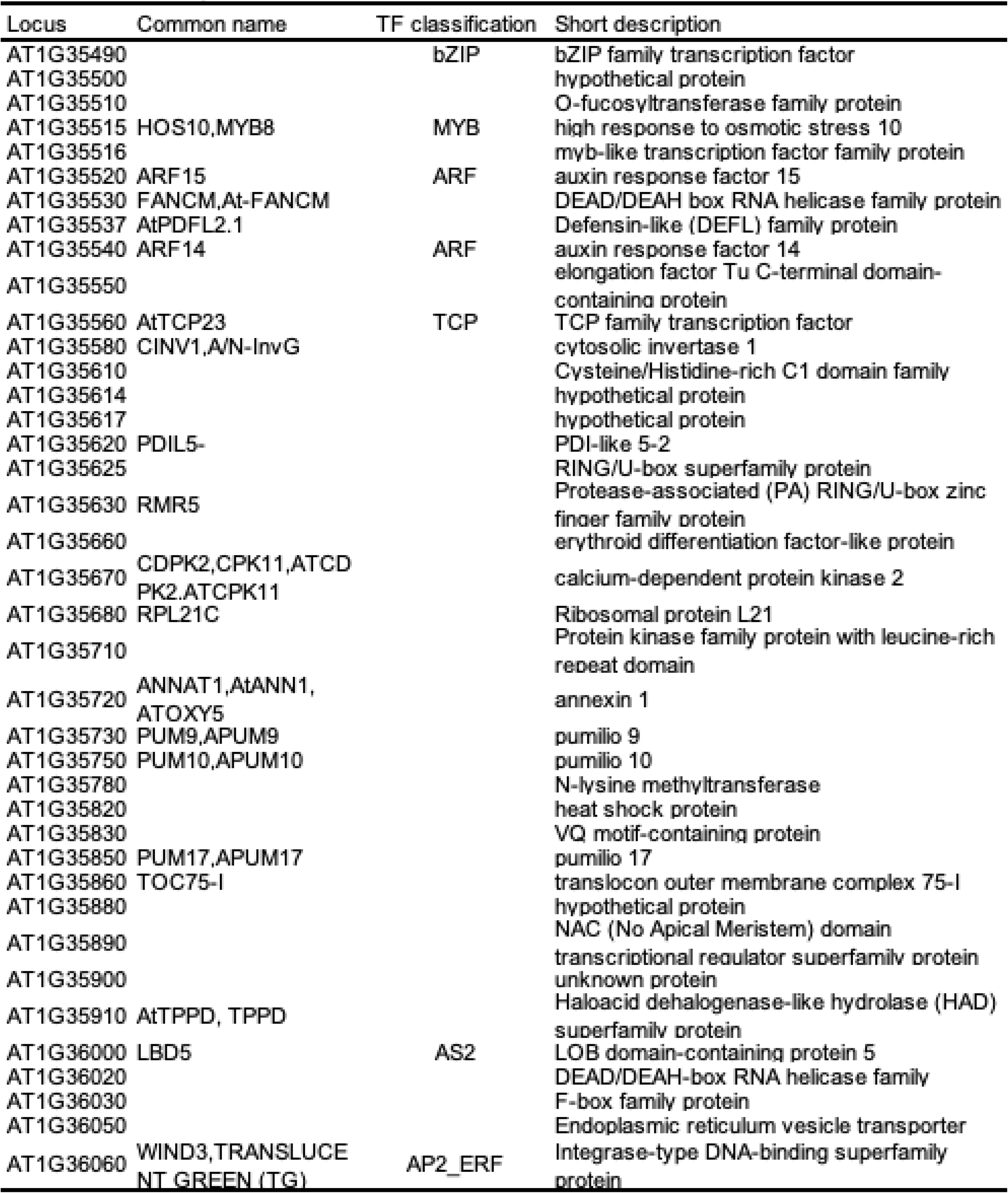
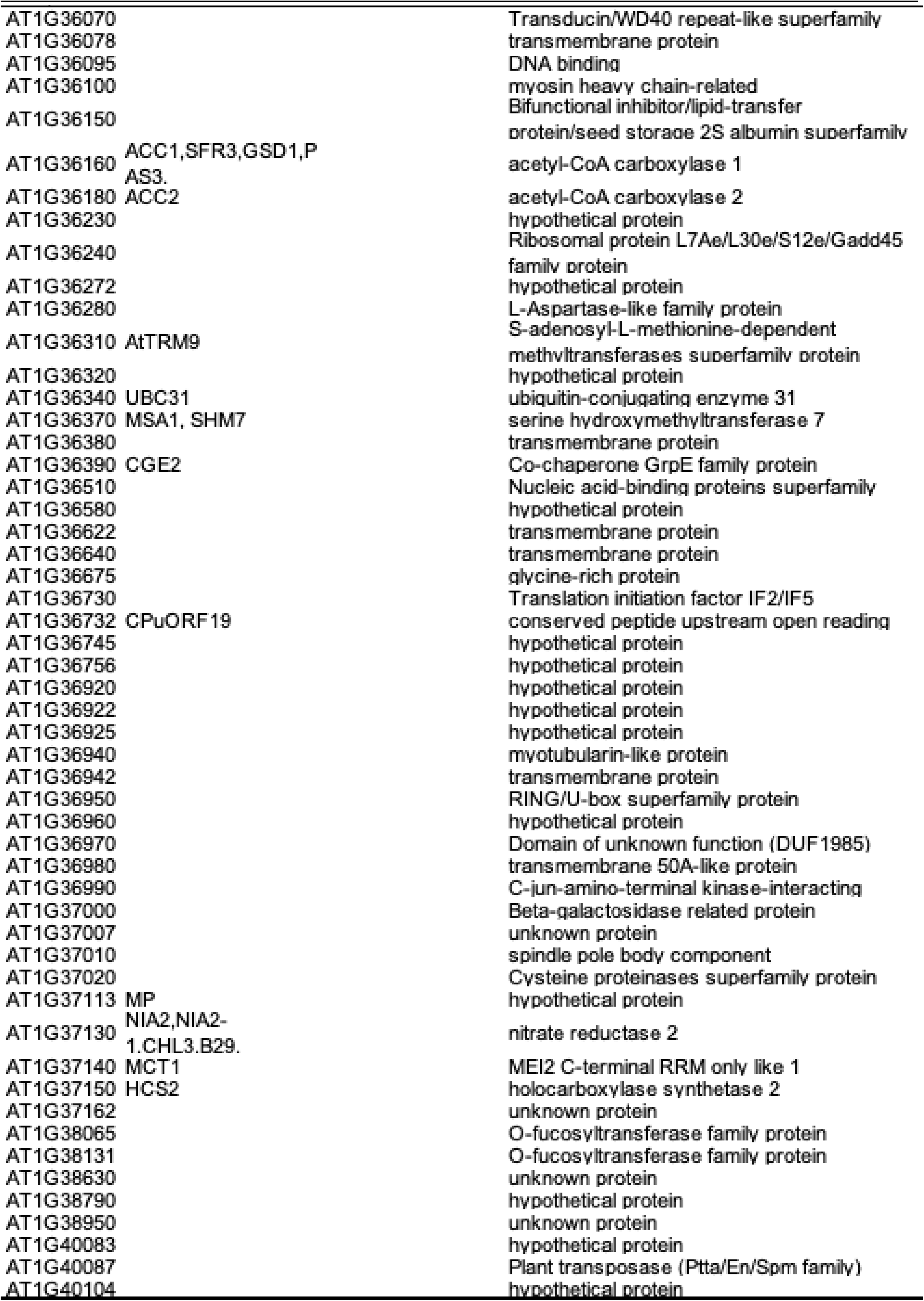
Deleted genes in Ar-7G-N2.

**Table S11.** Deleted genes in Ar-94-as1.

| Locus | Common name | TF classification | Short description |
| --- | --- | --- | --- |
| AT1G63160 | RFC2,EMB2811 |  | replication factor C 2 |
| AT1G63170 |  |  | Zinc finger, C3HC4 type (RING finger) family |
| AT1G63180 | UGE3 |  | UDP-D-glucose/UDP-D-galactose 4- |
| AT1G63190 |  |  | Cystatin/monellin superfamily protein |
| AT1G63200 |  |  | Cystatin/monellin superfamily protein |
| AT1G63205 |  |  | Cystatin/monellin superfamily protein |
| AT1G63206 |  |  | Cystatin/monellin superfamily protein |
| AT1G63210 | SPT6 |  | Transcription elongation factor Spt6 |
| AT1G63220 |  |  | Calcium-dependent lipid-binding (CaLB domain) family protein |
| AT1G63230 |  |  | Tetratricopeptide repeat (TPR)-like superfamily protein |
| AT1G63240 |  |  | hypothetical protein |
| AT1G63245 | CLE14 |  | CLAVATA3/ESR-RELATED 14 |
| AT1G63250 |  |  | DEA(D/H)-box RNA helicase family protein |
| AT1G63260 | TET10 |  | tetraspanin10 |
| AT1G63270 | AtCCMA,ABCI1, ATNAP10 |  | non-intrinsic ABC protein 10 |
| AT1G63280 |  |  | Serine protease inhibitor (SERPIN) family |
| AT1G63290 |  |  | Aldolase-type TIM barrel family protein |
| AT1G63295 |  |  | Remorin family protein |
| AT1G63300 |  |  | Myosin heavy chain-related protein |
| AT1G63310 |  |  | hypothetical protein |
| AT1G63320 |  |  | Pentatricopeptide repeat (PPR) superfamily |
| AT1G63330 |  |  | Pentatricopeptide repeat (PPR) superfamily |
| AT1G63340 |  |  | Flavin-containing monooxygenase family |

**Table S12.** Deleted genes in Ar-94-as1 (1)

| Locus | Common name | TF classification | Short description |
| --- | --- | --- | --- |
| AT3G30160 |  |  | transmembrane protein |
| AT3G30180 | BR6OX2,CYP85A2 |  | brassinosteroid-6-oxidase 2 |
| AT3G30200 |  |  | Plant transposase (Ptta/En/Spm family) |
| AT3G30210 | MYB121,ATMYB121 | MYB | myb domain protein 121 |
| AT3G30220 |  |  | hypothetical protein |
| AT3G30222 |  |  | transmembrane protein |
| AT3G30230 |  |  | myosin heavy chain-related |
| AT3G30235 |  |  | General transcription factor 2-related zinc finger protein |
| AT3G30247 |  |  | Protein of unknown function (DUF784) |
| AT3G30260 | AGL79 | MADS | AGAMOUS-like 79 |
| AT3G30280 |  |  | HXXXD-type acyl-transferase family protein |
| AT3G30290 | CYP702A8 |  | cytochrome P450, family 702, subfamily A, polypeptide 8 |
| AT3G30300 |  |  | O-fucosyltransferase family protein |
| AT3G30320 |  |  | hypothetical protein |
| AT3G30340 | UMAMIT32 |  | nodulin MtN21/EamA-like transporter family |
| AT3G30350 | RGF4,GLV3 |  | root meristem growth factor-like protein; root meristem growth factor 4 |
| AT3G30370 |  |  | MuDR family transposase |
| AT3G30380 |  |  | alpha/beta-Hydrolases superfamily protein |
| AT3G30383 |  |  | Protein of unknown function (DUF784) |
| AT3G30385 |  |  | Protein of unknown function (DUF784) |
| AT3G30387 |  |  | Protein of unknown function (DUF784) |
| AT3G30390 |  |  | Transmembrane amino acid transporter |
| AT3G30391 |  |  | unknown protein |
| AT3G30430 |  |  | F-box family protein |
| AT3G30456 |  |  | unknown protein |
| AT3G30460 |  |  | RING/U-box superfamily protein |
| AT3G30520 |  |  | heat shock protein |
| AT3G30530 | bZIP42,ATBZIP42 | bZIP | basic leucine-zipper 42 |
| AT3G30540 |  |  | Glycosyl hydrolase superfamily protein |
| AT3G30580 |  |  | hypothetical protein |
| AT3G30705 |  |  | transmembrane protein |
| AT3G30720 | QQS |  | qua-quine starch |
| AT3G30725 | GDU6,ATGDU6 |  | glutamine dumper 6 |
| AT3G30730 |  |  | hypothetical protein |

**Table S13.**
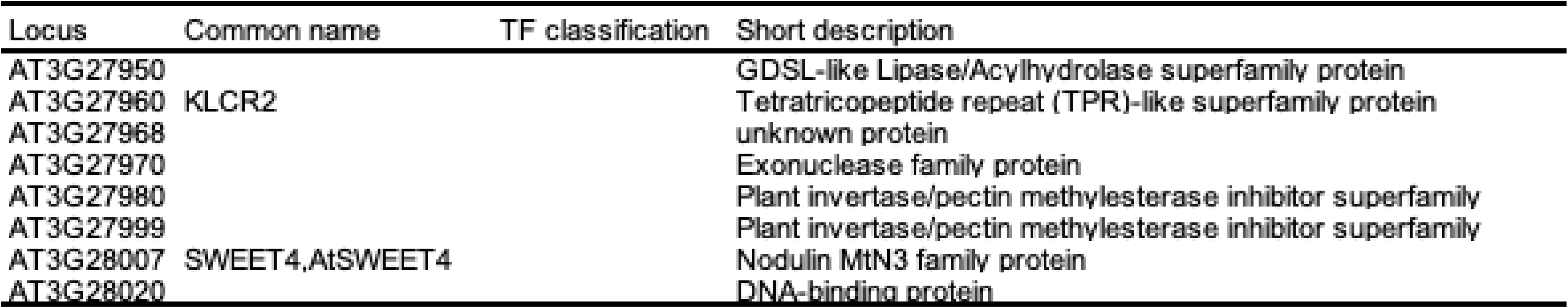
Deleted gene in C100-2^-N2.

**Table S14.**
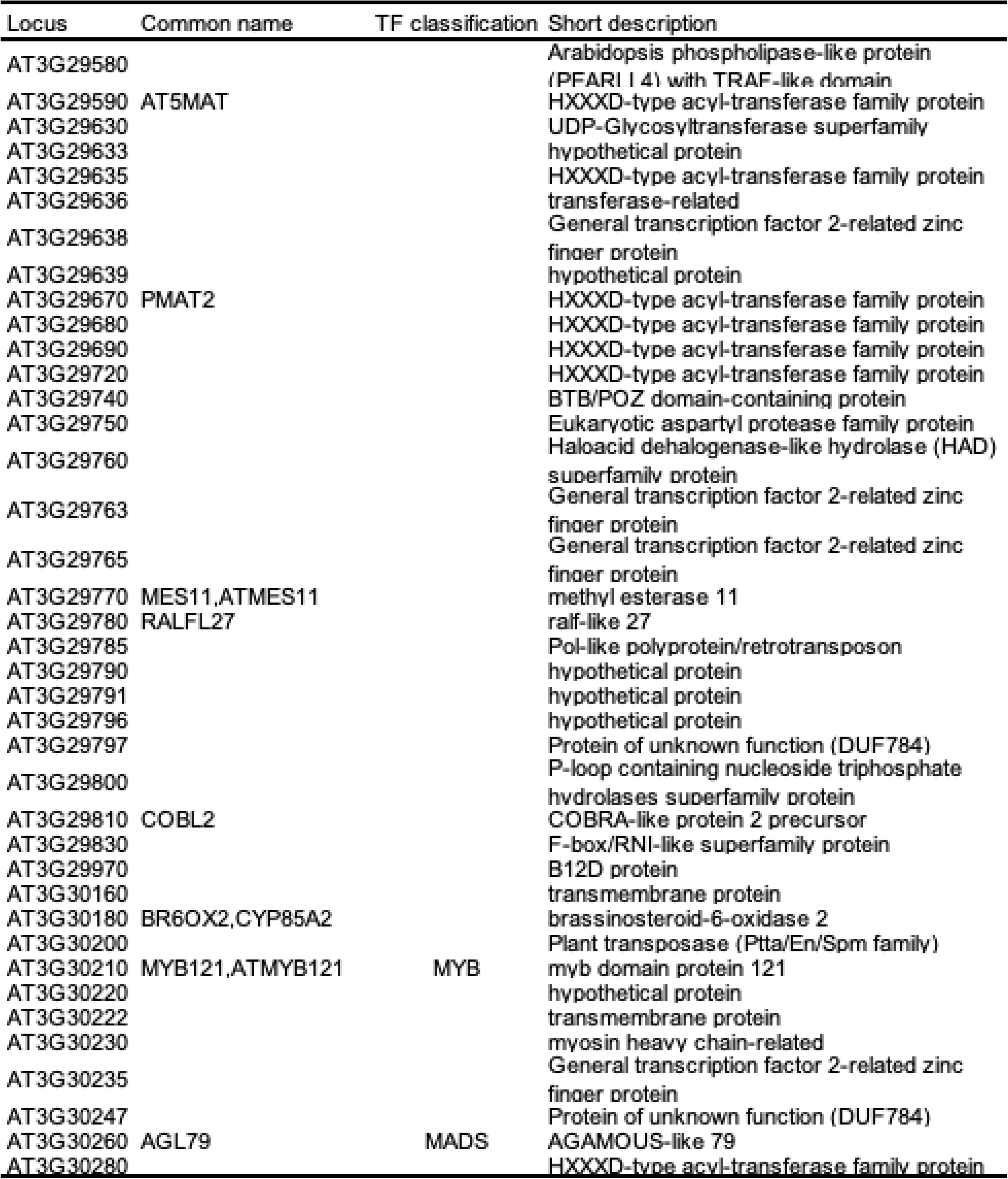
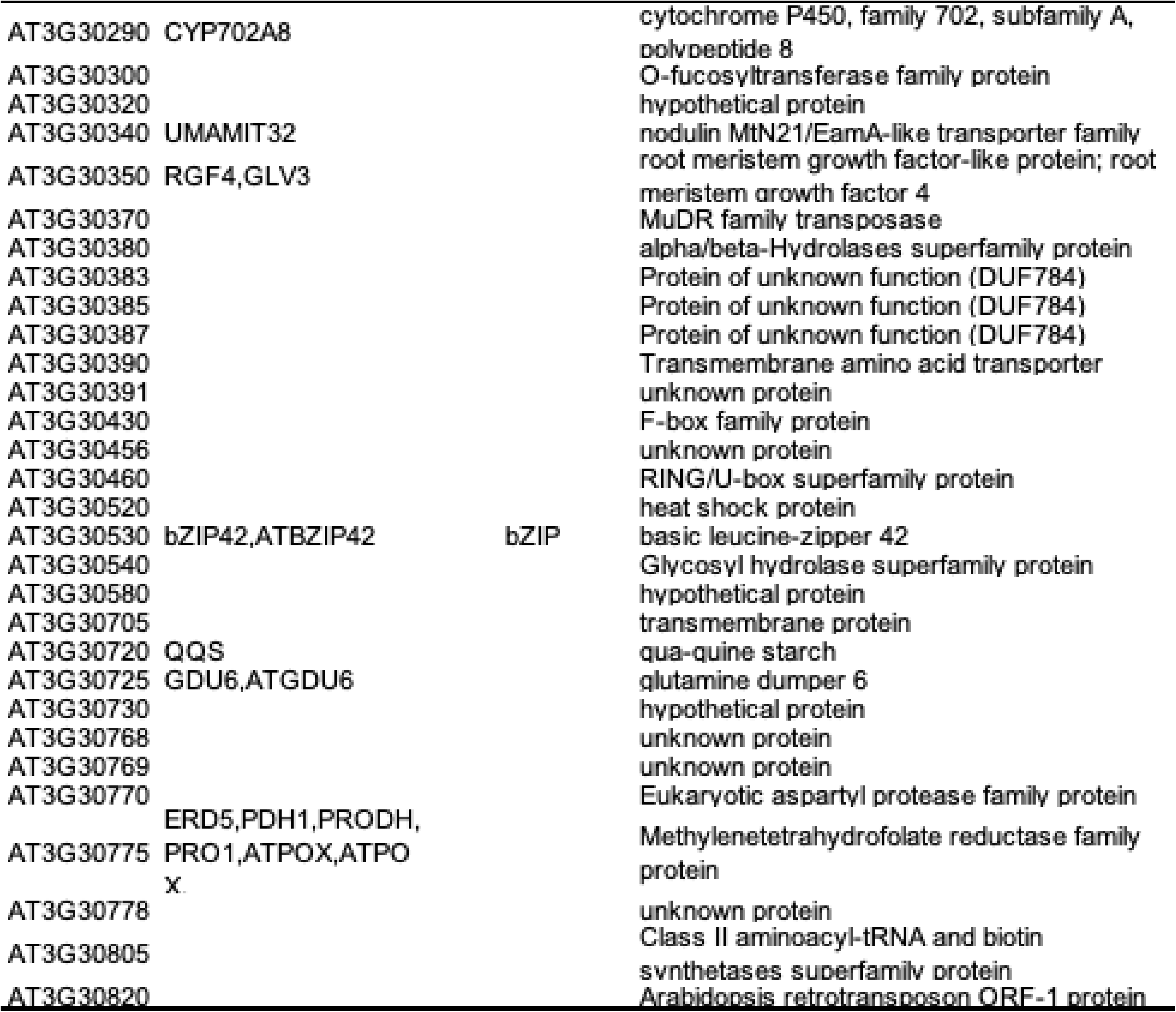
Deleted genes in C200-56-as4.

**Table S15.**
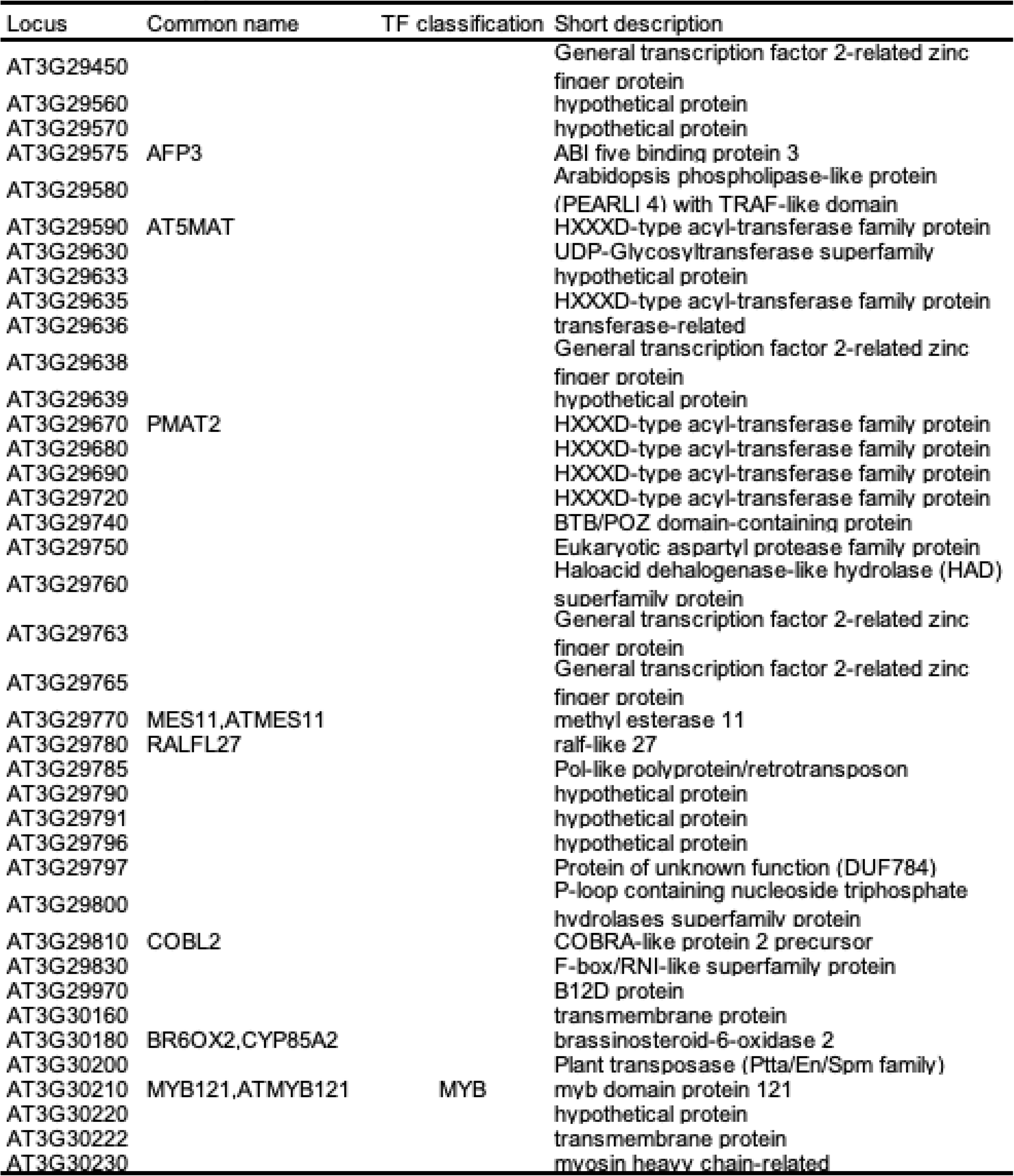
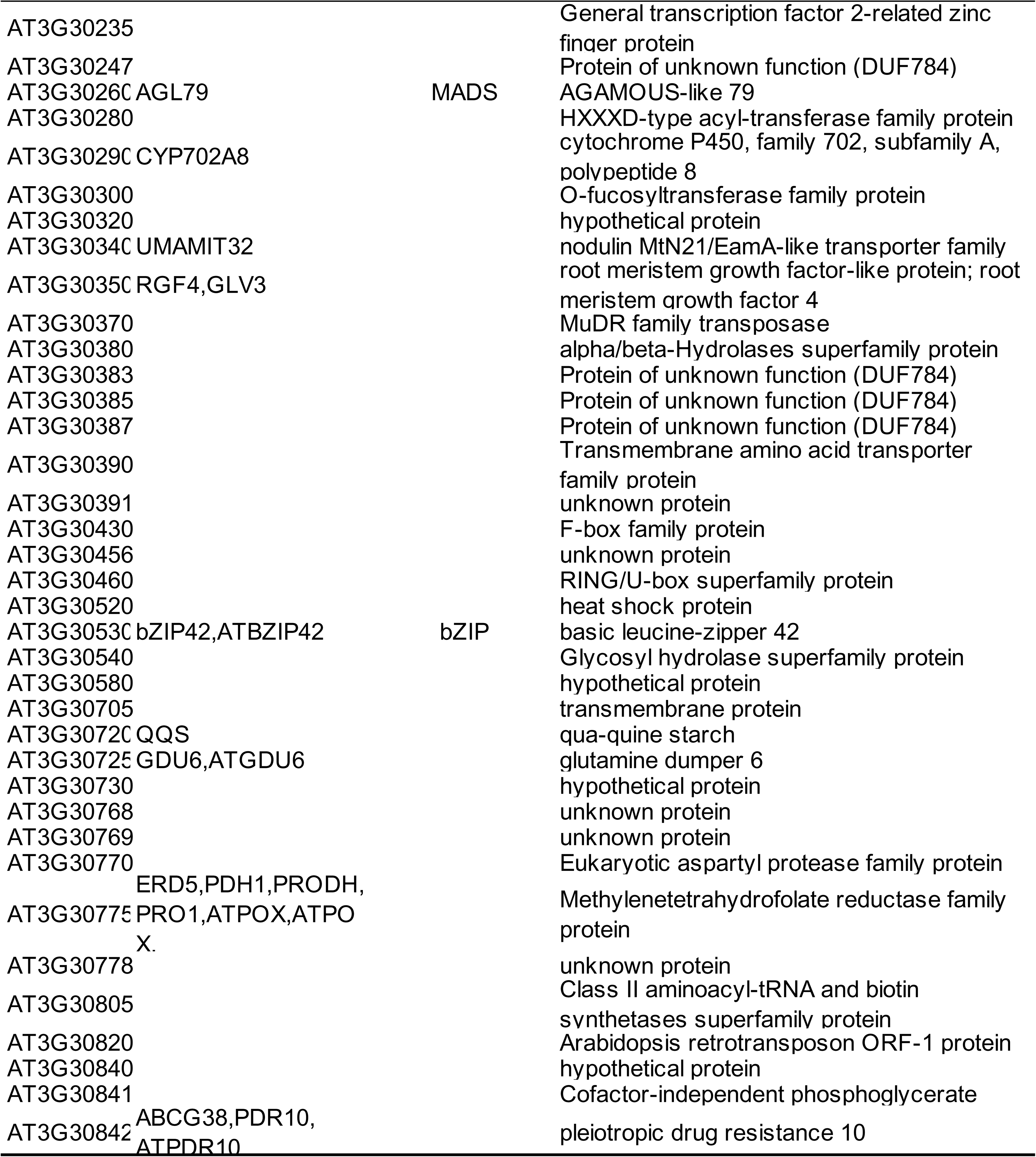
Deleted genes in C3O-144-as3.

**Table S16.**
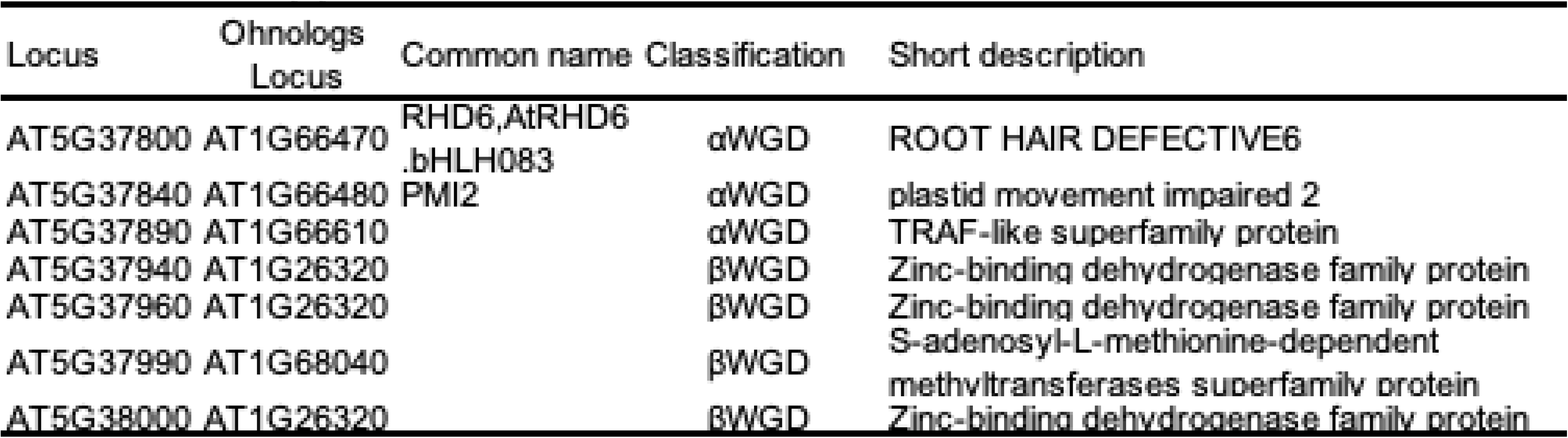
Oh nolog genes included in the deletion of AM1-N1.

**Table S17.** Oh nolog genes included in the deletion of Ar-47-N1

| Locus | Ohnologs<br>Locus | Common name | Classification | Short description |
| --- | --- | --- | --- | --- |
| AT3G46230 | AT5G59720 | HSP18.2,HSP<br>18.2.HSP18.1 | αWGD | heat shock protein 18.2 |
| AT3G46290 | AT5G5970 |  | αWGD | Protein kinase superfamily protein |
| AT3G46330 | AT5G5965<br>0 |  | αWGD | Leucine-rich repeat protein kinase<br>family protein |
| AT3G46420 | AT1G0755<br>0 |  | βWGD | Leucine-rich repeat protein kinase<br>family protein |
| AT3G46440 | AT5G5929 | UXS3,ATUXS3 | αWGD | UDP-glucuronic acid decarboxylase 3 |
|  | AT2G2875 |  | βWGD | UDP-D-glucuronate carboxy-lyase-related |
| AT3G46460 | AT5G5930 | UBC7,ATUBC7 | αWGD | ubiquitin carrier protein 7 |
| AT3G46510 | AT2G2883 | PUB12,AtPUB1 | βWGD | PLANT U-BOX 12 |
| AT3G46520 | AT5G5937 | ACT4 | αWGD | actin 4 |
| AT3G46580 | AT5G5938<br>0 | MBD6,ATMBD6 | αWGD | methyl-CPG-binding domain 6 |
| AT3G46590 | AT5G5943 | TRP1,ATTRP1 | αWGD | telomeric repeat binding protein 1 |
|  | AT1G0754 | TRFL2 | βWGD | TRF-like 2 |
| AT3G46600 | AT5G5945 | SCL11 | αWGD | GRAS family transcription factor |
|  | AT2G2906 |  | βWGD | GRAS family transcription factor |
| AT3G46620 | AT5G5955<br>0 | RDUF2,AtRDU | αWGD | zinc finger (C3HC4-type RING finger)<br>family protein |
| AT3G46640 | AT5G5957 | BOA | αWGD | Homeodomain-like superfamily protein |
| AT3G46650 | AT5G5959 | UGT76E2 | αWGD | UDP-glucosyl transferase 76E2 |
| AT3G46760 | AT5G5927<br>0 | LecRK-II.2 | αWGD | Concanavalin A-like lectin protein kinase<br>family protein |

**Table S18.**
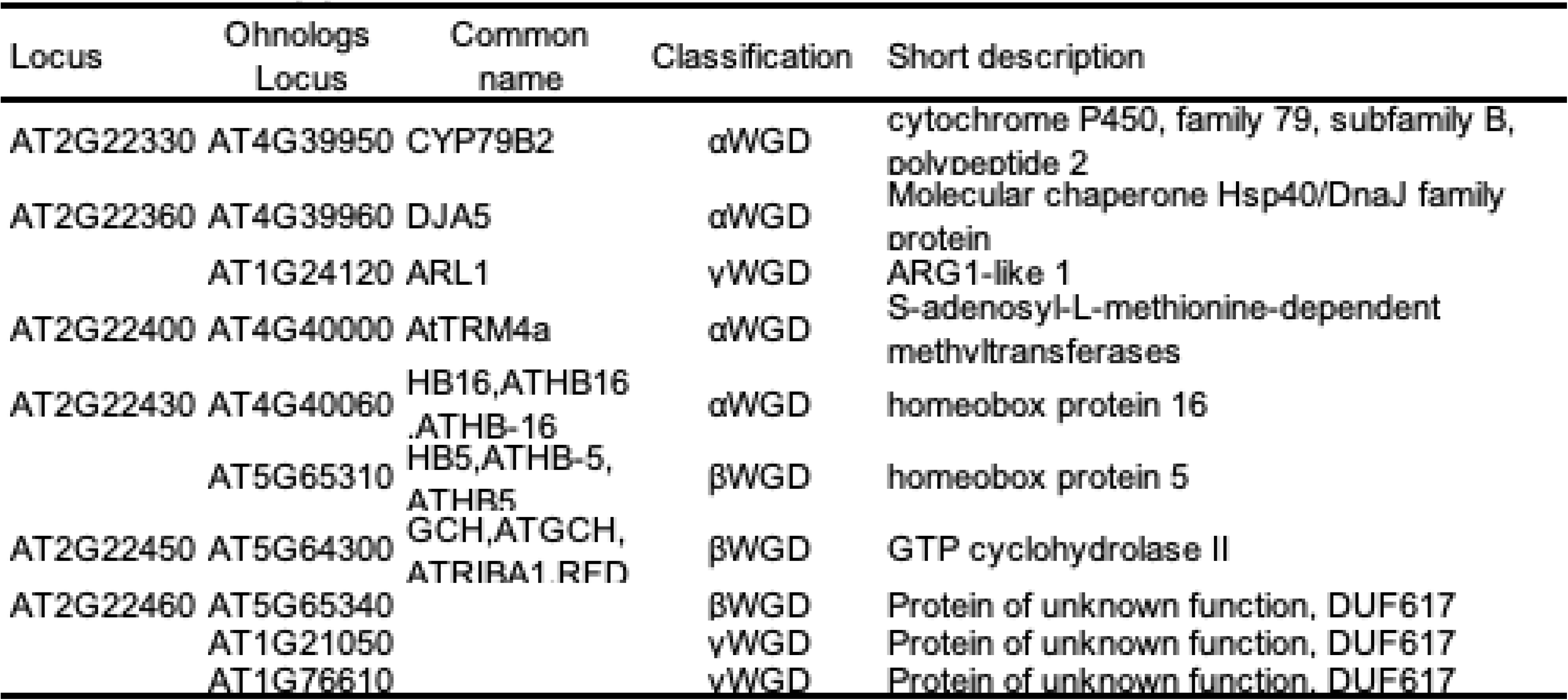
Oh nolog genes included in the deletion of Ar-57-as4.

**Table S19.**
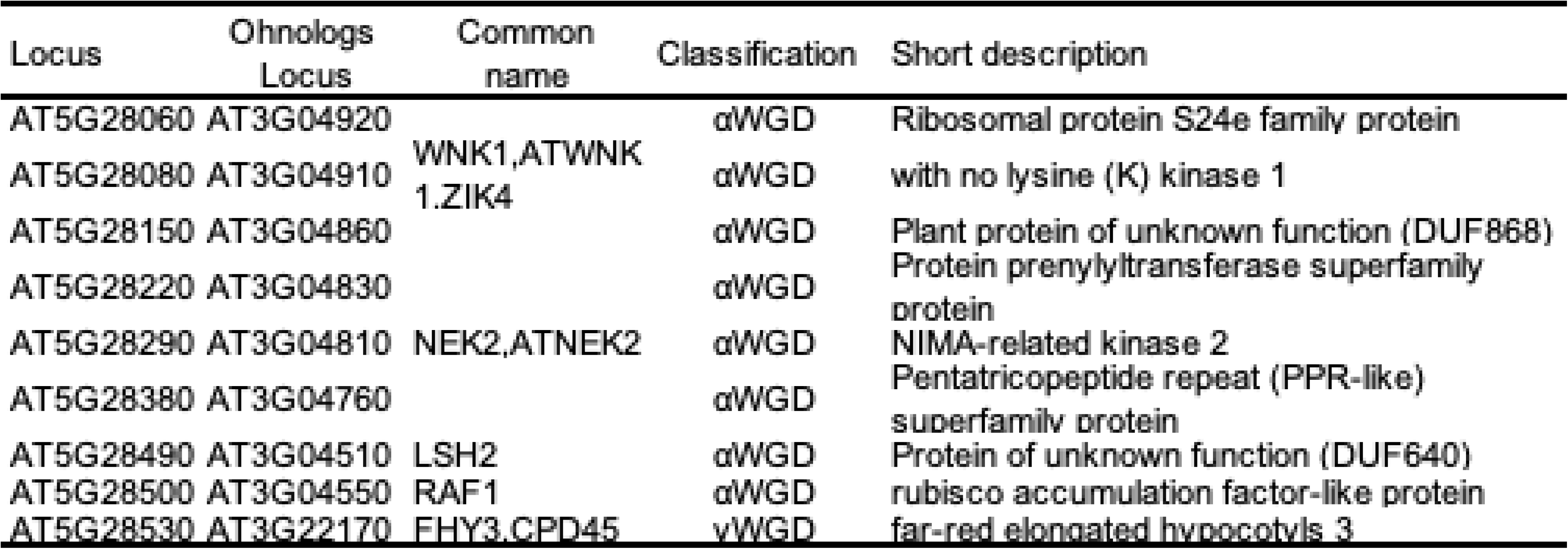
Oh nolog genes included in the deletion of Ar-S&-as6.

**Table S20.** Oh nolog genes included in the deletion of Ar-70-N2.

| Locus | Ohnologs<br>Locus | Common<br>name | Classification | Short description |
| --- | --- | --- | --- | --- |
| AT1G35515 | AT4G09460 | MYB6,ATMYB6,ATMYB8 | αWGD | myb domain protein 6 |
| AT1G35516 | AT1G22640 | MYB3,AtMYB3 | βWGD | myb domain protein 3 |
| AT1G35560 | AT1G72010 | AtTCP22,<br>TCP22 | βWGD | TCP family transcription factor |
| AT1G35580 | AT4G09510 | CINV2,A/N-Invl | αWGD | cytosolic invertase 2 |
| | AT1G22650 | A/N-InvD | βWGD | Plant neutral invertase family protein |
| AT1G35625 | AT4G09560 | RMR4, NHL22 | αWGD | Protease-associated (PA) RING/U-box zinc<br>finger |
| | AT1G71980 | RMR2 | βWGD | Protease-associated (PA) RING/U-box zinc<br>finger |
| AT1G35630 | AT1G71980 | RMR2 | βWGD | Protease-associated (PA) RING/U-box zinc<br>finger |
| AT1G35670 | AT4G09570 | CPK4,ATCPK4 | αWGD | calcium-dependent protein kinase 4 |

**Table S21.** Ohnolog genes included in the deletion of Ar-94-as1.

| Locus | Ohnologs<br>Locus | Common<br>name | Classification | Short description |
| --- | --- | --- | --- | --- |
| AT1G63170 | AT1G12760 | | αWGD | Zinc finger, C3HC4 type (RING finger)<br>family protein |
| | AT4G11680 | | βWGD | Zinc finger, C3HC4 type (RING finger)<br>family protein |
| AT1G63180 | AT1G12780 | UGE1,ATUGE1 | αWGD | UDP-D-glucose/UDP-D-galactose 4-epimerase |
| AT1G63300 | AT5G41140 | | αWGD | Myosin heavy chain-related protein |

**Table S22.** Oh nolog genes included in the deletion of C100-23-N2.

| Locus | Ohnologs<br>Locus | Common<br>name | Classification | Short description |
| --- | --- | --- | --- | --- |
| AT3G27950 | AT5G14450 | | βWGD | GDSL-like Lipase/Acylhydrolase<br>superfamily protein |
| | AT1G67830 | FXG1,ATFXG' | γWGD | alpha-fucosidase 1 |
| AT3G27970 | AT5G40310 | | αWGD | Exonuclease family protein |
| AT3G28007 | AT5G40260 | SWEET8.RPG | αWGD | Nodulin MtN3 family protein |

**Table S23.**
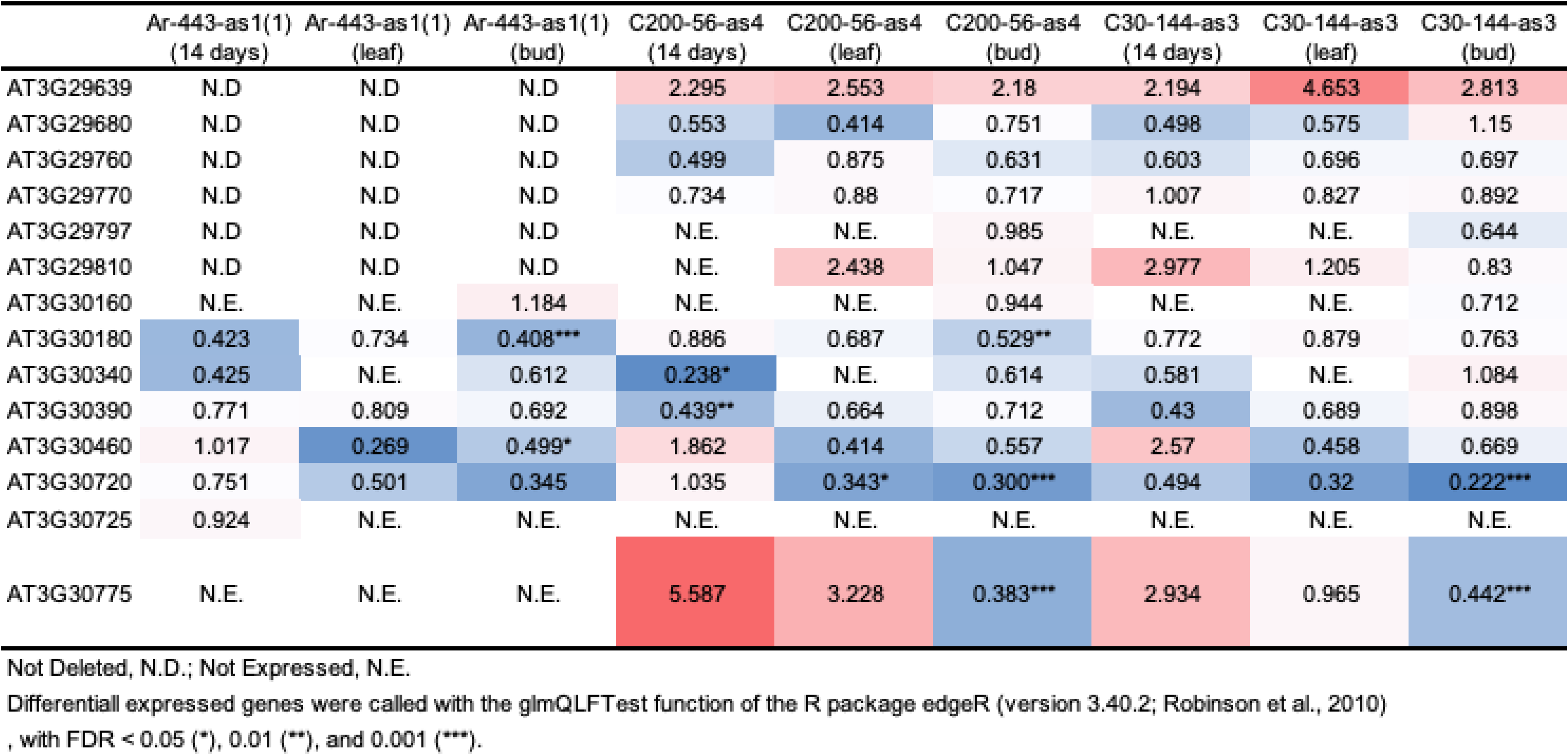
Fold change of gene expression of heterozygous ly deleted regions which were sheard among Ar-443-as1 (1), C200-56-as4, and C3O-144-asi mutants with respect to CobO.

|  | Ar-443-as1(1)<br>(14 days) | Ar-443-as1(1)<br>(leaf) | Ar-443-as1(1)<br>(bud) | C200-56-as4<br>(14 days) | C200-56-as4<br>(leaf) | C200-56-as4<br>(bud) | C30-144-as3<br>(14 days) | C30-144-as3<br>(leaf) | C30-144-as3<br>(bud) |
| --- | --- | --- | --- | --- | --- | --- | --- | --- | --- |
| AT3G29639 | N.D | N.D | N.D | 2.295 | 2.553 | 2.18 | 2.194 | 4.653 | 2.813 |
| AT3G29680 | N.D | N.D | N.D | 0.553 | 0.414 | 0.751 | 0.498 | 0.575 | 1.15 |
| AT3G29760 | N.D | N.D | N.D | 0.499 | 0.875 | 0.631 | 0.603 | 0.696 | 0.697 |
| AT3G29770 | N.D | N.D | N.D | 0.734 | 0.88 | 0.717 | 1.007 | 0.827 | 0.892 |
| AT3G29797 | N.D | N.D | N.D | N.E. | N.E. | 0.985 | N.E. | N.E. | 0.644 |
| AT3G29810 | N.D | N.D | N.D | N.E. | 2.438 | 1.047 | 2.977 | 1.205 | 0.83 |
| AT3G30160 | N.E. | N.E. | 1.184 | N.E. | N.E. | 0.944 | N.E. | N.E. | 0.712 |
| AT3G30180 | 0.423 | 0.734 | 0.408*** | 0.886 | 0.687 | 0.529** | 0.772 | 0.879 | 0.763 |
| AT3G30340 | 0.425 | N.E. | 0.612 | 0.238* | N.E. | 0.614 | 0.581 | N.E. | 1.084 |
| AT3G30390 | 0.771 | 0.809 | 0.692 | 0.439** | 0.664 | 0.712 | 0.43 | 0.689 | 0.898 |
| AT3G30460 | 1.017 | 0.269 | 0.499* | 1.862 | 0.414 | 0.557 | 2.57 | 0.458 | 0.669 |
| AT3G30720 | 0.751 | 0.501 | 0.345 | 1.035 | 0.343* | 0.300*** | 0.494 | 0.32 | 0.222*** |
| AT3G30725 | 0.924 | N.E. | N.E. | N.E. | N.E. | N.E. | N.E. | N.E. | N.E. |
| AT3G30775 | N.E. | N.E. | N.E. | 5.587 | 3.228 | 0.383*** | 2.934 | 0.965 | 0.442*** |

Not Deleted, N.D.; Not Expressed, N.E.
Differentiall expressed genes were called with the glmQLFTest function of the R package edgeR (version 3.40.2; Robinson et al., 2010) , with FDR < 0.05 (\*), 0.01 (\*\*), and 0.001 (\*\*\*).

